# A genetic model for *in vivo* proximity labeling of the mammalian secretome

**DOI:** 10.1101/2022.04.13.488228

**Authors:** Rui Yang, Amanda S. Meyer, Ilia A. Droujinine, Namrata D. Udeshi, Yanhui Hu, Jinjin Guo, Jill A. McMahon, Dominique K. Carey, Charles Xu, Qiao Fang, Jihui Sha, Shishang Qin, David Rocco, James Wohlschlegel, Alice Y. Ting, Steven A. Carr, Norbert Perrimon, Andrew P. McMahon

**Affiliations:** Department of Stem Cell Biology and Regenerative Medicine, University of Southern California, Los Angeles, CA, USA; Eli and Edythe Broad Center for Regenerative Medicine and Stem Cell Research, University of Southern California, Los Angeles, CA, USA; Department of Molecular Medicine, Scripps Research, La Jolla, CA, USA; Broad Institute of Harvard and MIT, Cambridge, MA, USA; Department of Genetics, Blavatnik Institute, Harvard Medical School, Boston, MA, USA; Department of Molecular Genetics, University of Toronto, Toronto, ON M5S 3E1, Canada; Department of Biological Chemistry, Geffen School of Medicine at UCLA, University of California, Los Angeles, Los Angeles, CA; BIOPIC, Beijing Advanced Innovation Center for Genomics, School of Life Sciences, Peking University, Beijing, China; Chan Zuckerberg Biohub, San Francisco, CA, USA; Departments of Genetics, Biology, and Chemistry, Stanford University, Stanford, CA, USA; Howard Hughes Medical Institute, Boston, MA, USA

**Keywords:** proximity-labeling, biotin, BirA, TurboID, secretome, inter-organ communication, serum proteins

## Abstract

Organ functions are highly specialized and interdependent. Secreted factors regulate organ development and mediate homeostasis through serum trafficking and inter-organ communication. Enzyme-catalyzed proximity labeling enables the identification of proteins within a specific cellular compartment. Here, we report a *BirA*G3* mouse strain that enables CRE-dependent promiscuous biotinylation of proteins trafficking through the endoplasmic reticulum. When broadly activated throughout the mouse, widespread labeling of proteins was observed within the secretory pathway. Streptavidin affinity purification and peptide mapping by quantitative mass spectrometry (MS) proteomics revealed organ-specific secretory profiles and serum trafficking. As expected, secretory proteomes were highly enriched for signal peptide-containing proteins, highlighting both conventional and non-conventional secretory processes, and ectodomain shedding. Lower-abundance proteins with hormone-like properties were recovered and validated using orthogonal approaches. Hepatocyte-specific activation of BirA*G3 highlighted liver-specific biotinylated secretome profiles. The BirA*G3 mouse model demonstrates enhanced labeling efficiency and tissue specificity over viral transduction approaches and will facilitate a deeper understanding of secretory protein interplay in development, and healthy and diseased adult states.

## Introduction

Protein secretion plays a critical role in coordinating local and systemic cellular responses in development, homeostasis, and disease^1–4^. Multi-organ failure suggests an aberrant organ–organ crosstalk resulting in linked organ pathology such as pulmonary–renal syndromes^5, 6^, with an increased risk of sequential organ failure and morbidity^7^. Secreted proteins may be identified using liquid chromatography-tandem mass spectrometry (LC-MS/MS) proteomics of serum^8^. However, it is challenging to identify low abundance proteins and difficult to track the organs of origin and ultimate destination of protein interactions^9, 10^.

Analysis of cell secretomes has benefited from enzyme-catalyzed proximity labeling approaches such as BioID^11^ and TurboID^12^. In these, the activity of a promiscuous biotin-ligase in the cellular secretory pathway biotinylates resident and secreted proteins, which can then be detected by affinity enrichment and quantitative MS^11, 12^. These labeling methods allow sensitive and stable detection of endogenous secreted proteome in live cells, including fly (*Drosophila melanogaster*)^12, 13^ and worm (*Caenorhabditis elegans*) models^12^, mouse tumor transplants^13^ and specific mammalian target tissues through viral directed gene delivery^14–16^. These studies have provided new insight into tissue secretomes and inter-organ communication^14–19^.

To overcome the limitations of viral-mediated approaches for systematic temporal and spatial analysis of mammalian cell secretomes *in vivo*, we generated and validated a mouse model system. In the secretome reporter strain, DNA sequences encoding an endoplasmic reticulum directed promiscuous biotin ligase, BirA*G3^12^, were inserted into the ubiquitously expressed Rosa26 locus^13^. BirA*G3 is a precursor to TurboID, generated in the directed evolution of *E. coli* BirA, that is more active than TurboID and has higher affinity for biotin^12^. Conditional (CRE recombinase- and exogenous biotin-dependent) BirA*G3 activity resulted in rapid biotinylation of proteins trafficking through the secretory pathway and permitted the analysis of cellular secretomes through streptavidin affinity purification and quantitative mass spectrometry proteomics ^12, 13^.

## Results

### Activation of BirA*G3 in *Sox2-BirA*G3* mice

We previously reported a Cre-inducible *BirA*G3* cassette, inserted in the *Rosa26 (R26)* “safe-harbor” locus in mouse embryo stem cells (mESCs) (Fig. 1A)^13^. Here, we derived mouse strains, from three independently targeted mESCs containing the Cre-inducible *BirA*G3* cassette (A11, B1, and C2). As expected, no differences were observed comparing the three lines consequently; we refer to data as if from a single line and we chose the A11 line for more extensive characterization.

**Fig. 1.**
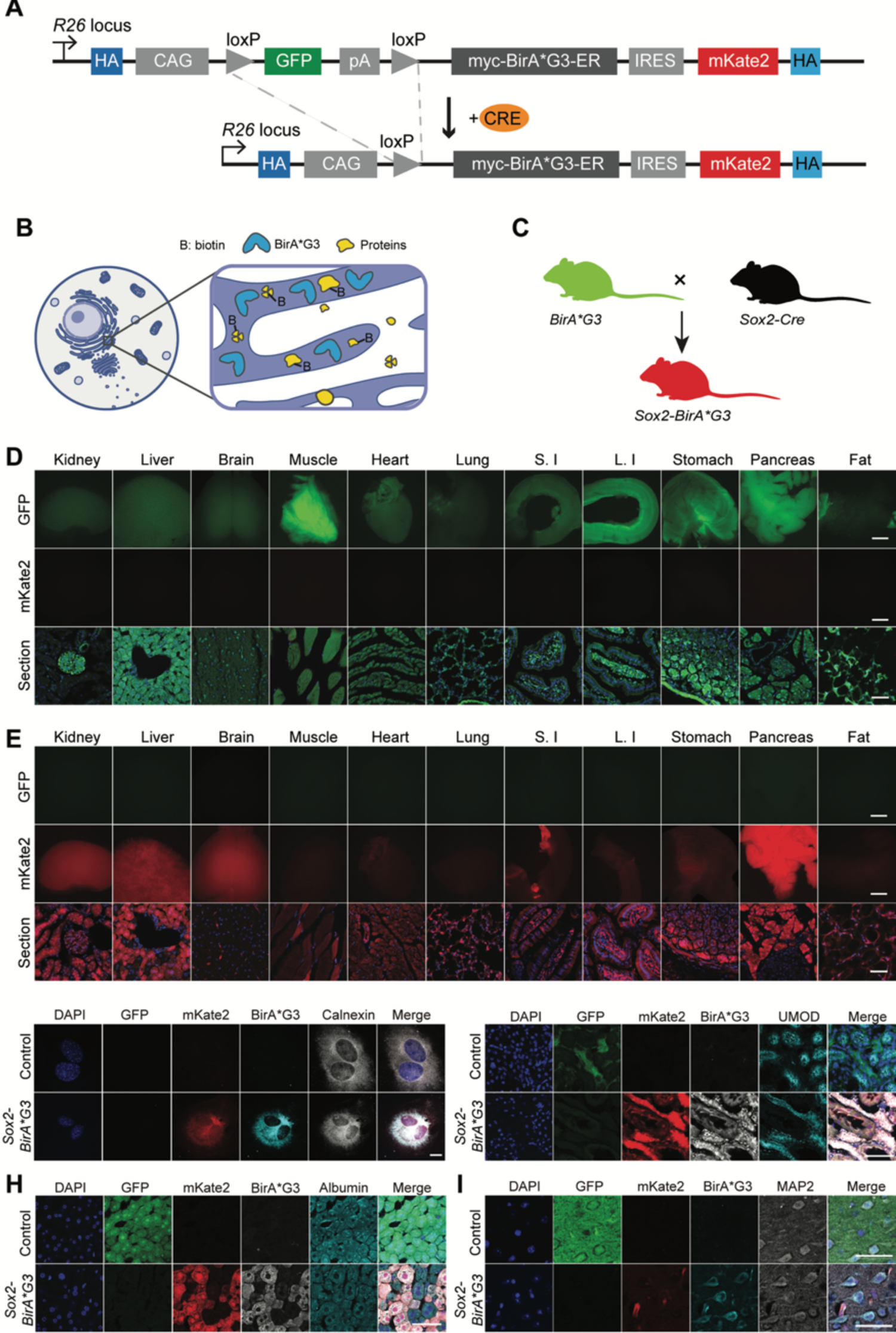
Generation and characterization of *Sox2-BirA*G3* mice. (A) Schematic diagram shows CRE-mediated excision at loxP sites removes the GFP cassette resulting in production of BirA*G3-ER and mKate2 fluorescent protein. Adapted from Droujinine et al. 2021. (B) Schematic diagram shows that proteins that reside or travel through ER would be biotinylated by BirA*G3. (C) Schematic diagram of mouse mating to generate *Sox2-BirA*G3* mice. All cells in control mice are expected to express GFP. All cells in *Sox2-BirA*G3* mice are expected to express mKate2. Adapted from Droujinine et al. 2021. (D) Native GFP and native mKate2 fluorescence in whole-mount organs (Scale Bar: 2mm) and tissue sections (Scale Bar: 50μm) of control mice. S.I: small intestine. L.I: large intestine. (E) Native GFP and native mKate2 fluorescence in whole-mount organs (Scale Bar: 2mm) and tissue sections (Scale Bar: 50μm) of *Sox2-BirA*G3* mice. (F) Immunofluorescence staining shows expression of native GFP, native mKate2, BirA*G3, and ER marker Calnexin in the cell culture of isolated mouse embryonic fibroblast (MEFs) from *Sox2-BirA*G3* and control mice. Scale Bar: 25μm. (G) Immunofluorescence staining shows expression of native GFP, native mKate2, BirA*G3, and UMOD, which marks the thick ascending limb of Henle’s loop (TALH), in the kidney sections of *Sox2-BirA*G3* and control mice. Scale Bar: 50μm. (H) Immunofluorescence staining shows expression of native GFP, native mKate2, BirA*G3, and Albumin, a hepatocyte marker, in the liver sections of *Sox2-BirA*G3* and control mice. Scale Bar: 50μm. (I) Immunofluorescence staining shows expression of native GFP, native mKate2, BirA*G3, and MAP2, a cortical neuron-specific protein, in the brain sections of *Sox2-BirA*G3* and control mice. Scale Bar: 50μm. Unlike BirA*G3, mKate2 expression is not restricted to the ER, which may result in varied signal intensity due to differences in cell morphology.

The modified Rosa26 locus was designed to drive ubiquitous GFP expression downstream of a *CAGGS* (*β-actin/CMV*) regulatory sequence. GFP transcription blocks downstream expression of a *BirA*G3* cassette and mKate2 reporter (Fig. 1A)^13^. BirA*G3 encodes a promiscuously active biotin ligase selected by protein evolution^12^ with a signal peptide and ER retention signal, designed to target and retain BirA*G3 within the cell’s secretory pathway (Fig. 1B). mKate2 encodes a monomeric, photostable, pH resistant, low toxicity, bright far-red fluorescent protein designed as a cellular indicator of cells producing BirA*G3 and activating biotinylation of secretory pathway proteins on biotin administration (Fig. 1B)^20–22^. CRE recombination at loxP sites flanking the GFP cassette enables the tissue and time-dependent expression of BirA*G3 and mKate2^13^.

To initially examine BirA*G3 activity body-wide to broadly calibrate the model, we crossed the *R26 BirA*G3* mice to the *Sox2-Cre* strain which results in recombination and predicted activation of BirA*G3 and mKate2 in the epiblast, and thereafter, in all cell types of the conceptus^23^. As a consequence, cells with the unrecombined BirA*G3 allele will be green (GFP+), while those undergoing CRE-mediated recombination in *Sox2-Cre; BirA*G3* (abbreviated as *Sox2-BirA*G3*) mice will be red (mKate2+) and BirA*G3 positive (Fig. 1C).

Viability and fertility were normal in *Sox2-BirA*G3* mice. As predicted, whole-mount images of selected organs from control mice (*BirA*G3/+* unless otherwise stated) were GFP+/mKate2-, with variable GFP intensity across organs (Fig. 1D), while *Sox2-BirA*G3* mice were GFP-/mKate2+, demonstrating excision of the GFP cassette and activation of mKate2 reporter (Fig. 1E). High-magnification confocal images showed that BirA*G3 extensively colocalized with the ER resident protein calnexin (Fig. 1F; Extended Fig. 1A), consistent with BirA*G3 localization to the ER. Focusing on the kidney, liver, and brain, we next sought to determine the distribution of BirA*G3 using representative cell markers, highlighting key cell populations.

In the *Sox2-BirA*G3* kidney, mKate2 signal and BirA*G3 were present throughout the nephron and collecting system highlighted by co-localization with different cell markers (Fig. 1G; Extended Fig. 1B-C; Supplementary Fig. 1A-B). In *Sox2-BirA*G3* liver, mKate2 signal and BirA*G3 were present in hepatocytes (Albumin+), though the distribution was patchy (Fig. 1H) and neighboring cholangiocytes (CK19+) showed much lower levels of reporter and BirA*G3 (Extended Fig. 1D). In *Sox2-BirA*G3* brain, mKate2 signal and BirA*G3 expression was present in neurons (MAP2+ representative cortical neurons), astrocytes (GFAP+), and microglia (IBA1+) (Fig. 1I; Extended Fig. 1E-F; Supplementary Fig. 1C). Taken together, the data indicate a broad cell and tissue distribution for BirA*G3 and mKate2, though levels vary significantly depending on the cell type.

To compare BirA*G3 and the mKate2 reporter with another *CAGGS* (*β-actin/CMV)* driven reporters targeted to a similar position with the same transcriptional orientation in the *R26* locus, we crossed *Sox2-Cre* mice with the widely used R26 TdTomato reporter mouse strain^24^. We observed homogeneous production of TdTomato in *Sox2-cre; TdTomato* (*Sox2-TdTomato*) tissues (Extended Fig. 2A-B) suggesting that uneven BirA*G3/mKate2 in *Sox2-BirA*G3* mice is specific to the BirA*G3 targeted locus.

To specifically examine the patchy hepatocyte distribution, mKate2^low^ and mKate2^high^ hepatocytes were sorted from *Sox2-BirA*G3* liver and analyzed by qPCR (Extended Fig. 2C). No *BirA*G3* mRNA expression was detected in control liver as expected (Extended Fig. 2D). In *Sox2-BirA*G3* liver, *BirA*G3* mRNA was present in both mKate2^low^ and mKate2^high^ cells, though *BirA*G3* mRNA levels were also lower in mKate2^low^ cells (Extended Fig. 2D). In contrast, albumin mRNA expression was comparable in mKate2^low^ and mKate2^high^ populations (Extended Fig. 2E). The lower *BirA*G3* mRNA expression was consistent with lower BirA*G3 protein level in mKate2^low^ hepatocytes (Extended Fig. 2F-H). Protein biotinylation is still observed in mKate2^low^ cells, although at lower levels (Extended Fig. 2F-H), Thus, a mosaic reduction in transcriptional activity of the BirA*G3 locus likely underlies variable levels of mKate2 and BirA*G3 in hepatocytes.

**Fig. 2.**
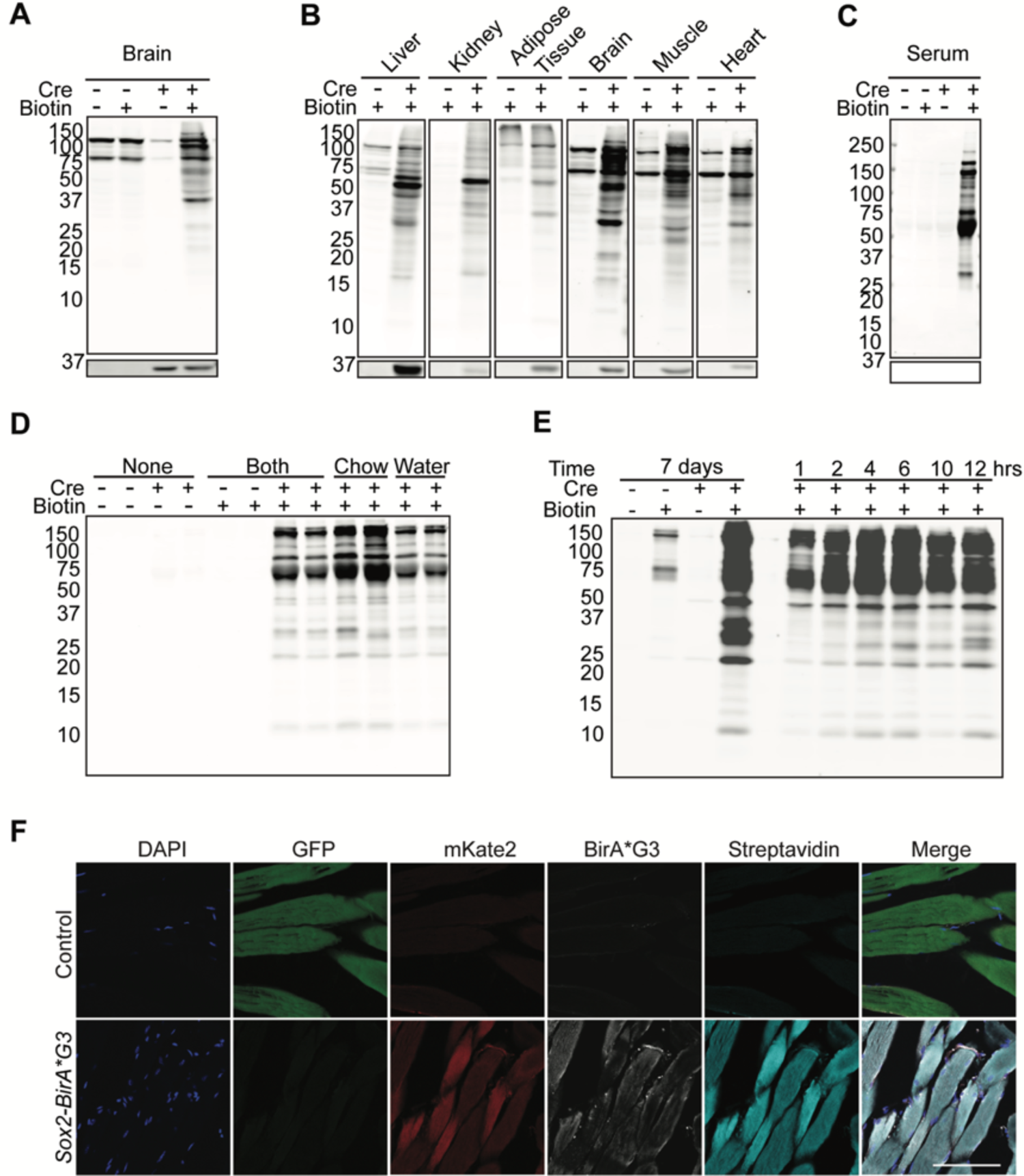
Analysis of biotinylated proteins in *Sox2-BirA*G3* and control mice. (A) Western blotting of protein lysates from brain in *Sox2-BirA*G3* mice (CRE+) compared to control mice (CRE-) with or without biotin chow administration for 5 days. Upper: Streptavidin labeling. Lower: BirA*G3 (∼35kDa). Each lane is a biological replicate from individual mice (n=2/genotype). (B) Western blotting of protein lysates from selective tissues in *Sox2-BirA*G3* mice compared to control mice. Note: due to varied streptavidin intensity by tissue, brain, heart, and muscle streptavidin westerns are shown with increased signal intensity. Upper: Streptavidin labeling. Lower: BirA*G3 (∼35kDa). Each lane is a biological replicate from individual mice (n=1/genotype). (C) Western blotting of serum total protein in *Sox2-BirA*G3* mice compared to control mice (CRE-) with or without biotin chow administration for 5 days. Upper: Streptavidin labeling. Lower: BirA*G3 (∼35kDa). Each lane is a biological replicate from individual mice (n=2/genotype). (D) Streptavidin labeling of biotinylated proteins in total serum from *Sox2-BirA*G3* and control mice administered with regular chow, biotin chow, biotin water, or biotin chow and water for 7 days. Each lane is a biological replicate from individual mice (n=2/biotin treatment). (E) Streptavidin labeling of affinity purified biotinylated proteins in serum from *Sox2-BirA*G3* and control mice given biotin chow for 7 days (left) or given biotin by subcutaneous injection and water (5mM, pH 7.4) after the injection until collection (right). (F) Immunofluorescence images of native GFP, native mKate2, BirA*G3 staining, streptavidin staining in cryo-sectioned muscle tissues from *Sox2-BirA*G3* and control mice. Scale Bar: 50μm.

To determine whether down-regulation relates to the duration of allele activation, we examined BirA*G3 in *Sox2-BirA*G3* and control pups 10 days after birth. Interestingly, at this early time point, BirA*G3 and mKate2 show a relatively homogenous distribution in hepatocytes (Extended Fig. 3A-B). While the molecular underpinnings for this observation are unclear, most cell/tissue types were not affected and inducible CRE strains can potentially overcome possible selection for allele silencing. Indeed, we observed homogenous BirA*G3 in GFP-hepatocytes of adult *CAGGCre-ER^TM^; BirA*G3* (*CAGG-BirA*G3*) mice following tamoxifen injection to induce broad, mosaic excision of the GFP cassette in the adult animal (Extended Fig. 3C-D).

### Protein Proximity labeling in *Sox2-BirA*G3* mice

Next, we characterized biotinylation of proteins in *Sox2-BirA*G3* and control mice. Mice were fed biotin in chow (2,000 ppm biotin) for 5 days *ad libitum.* Brain samples were collected and analyzed by western blotting to detect biotinylated proteins through streptavidin-IRDye 800CW binding. Prominent biotin/BirA*G3-dependent labeling was observed for a broad range of protein species (Fig. 2A). Additionally, we observed biotin/ BirA*G3 independent-labeling of two prominent proteins in control and experimental brain samples (Fig. 2A) which most likely represent cytoplasmic biotin conjugates with pyruvate carboxylase (∼130kDa) and methylcrotonyl-CoA carboxylase/propionyl-CoA carboxylase (∼75kDa)^14^(Fig. 2A; Supplementary Fig. 2A).

Examining a broad range of total protein lysates from a range of organs showed strong evidence for BirA*G3-dependent biotinylation of target tissues (Fig. 2B; Supplementary Fig. 2B). Notably, each tissue displayed a unique biotinylation pattern (Fig. 2B), suggesting diverse secretomes across tissues. Further, analysis of serum detected a robust BirA*G3-dependent biotinylated protein signature (Fig. 2C; Supplementary Fig. 2C). Comparable serum labeling was obtained through multiday (7 days) labeling with biotin addition to either water (5mM) or chow (2,000ppm) (Fig. 2D; Supplementary Fig. 2D, E). Subcutaneous injection of biotin (180mM, 100 μL) resulted in significant labeling of serum proteins within 1 hour of injection with an increased recovery of proteins up to 12 hours (Fig. 2E; Supplementary Fig. 2F). These results highlight the rapid kinetics for secretion of biotinylated proteins though comparison with protein products observed with continuous 7-day labeling in chow demonstrates enhanced labeling of lower molecular weight protein species (Figure 2E).

The accumulation of biotinylated proteins could also be visualized by streptavidin-dependent immunofluorescence directly in sections of selective tissues. Abundant biotinylated proteins were observed in *Sox2-BirA*G3* muscle, heart, brain, compared to their control counterparts (Fig. 2F; Supplementary Fig. 3A-B), consistent with CRE-dependent biotinylation. For the liver and kidney, endogenous biotin stores result in equivalent strong streptavidin signals in both *Sox2-BirA*G3* and control mice (Supplementary Fig. 2C-D), even though western analysis shows Sox2-BirA*\*G3* - dependent labeling of proteins (Fig. 2B).

To look for detrimental effects of BirA*G3 in *Sox2-*BirA*\*G3* mice, we performed hematoxylin and eosin staining and bulk mRNA-sequencing. No pathological changes were detected by histology comparing *Sox2-BirA*G3* and control mice fed biotin chow for 7 days (Extended Fig. 4A). Principal component analysis (PCA) of mRNA-sequencing comparing liver, brain, and kidney between control (n=2 per tissue) and *Sox2-BirA*G3* (n=2 per tissue) mice showed a tight clustering by tissue (Extended Fig. 4B). Differential gene expression analysis of all *Sox2-BirA*G3* samples compared to all control samples showed a total of only four differentially expressed genes (DEGs) (Extended Fig. 4C). Furthermore, analysis of ER stress, unfolded protein response (UPR), and cell death markers showed no difference (adj. p-value > 0.05) at the transcript level between *Sox2-BirA*G3* and control samples (Extended Fig. 4D). Additionally, western blotting showed no indication of ER stress or UPR in *Sox2-BirA*G3* and control livers compared to ER stress induced (tunicamycin treated) MEFs (Extended Fig. 4E-J).

### MS-based tissue proteomics of biotinylated proteins

As a prelude to MS analyses of each of the three tissues (liver, brain, and kidney), affinity purified biotinylated proteins were visualized by both streptavidin western analysis and silver stain. The biotinylation banding patterns were comparable to those observed in total protein lysates of corresponding samples, suggesting efficient and unbiased enrichment of biotinylated proteins (Extended Fig. 5A-D). Silver stain showed specific bands in *Sox2*-*BirA*G3* samples, despite a strong background of non-specific binding of unlabeled proteins to streptavidin conjugated beads in all three tissues (Extended Fig. 5E-H).

The biotinylated proteomes of the three tissues were defined by quantitative TMT-based LC-MS/MS following streptavidin bead enrichment from *Sox2-BirA*G3* and control liver, brain, and kidney. Of the thousands of proteins detected and quantified in the *Sox2-BirA*G3* tissues, several hundred proteins were found to be significantly enriched in liver (n=189), brain (n=200), kidney (n=578) compared to their control counterparts (log_2_ fold change (FC)>1.0 and adj. p-value<0.05) (Fig. 3A-B; Extended Fig. 6A-C; Supplementary Data 1-3). Principal component analysis (PCA) demonstrated tight grouping of biological replicates within the *Sox2-BirA*G3* group, indicating similar proteomic profiles among these samples (Supplementary Fig. 4A-C). Relative abundance of representative signature proteins for each tissue is shown in in Fig. 3C.

**Fig. 3.**
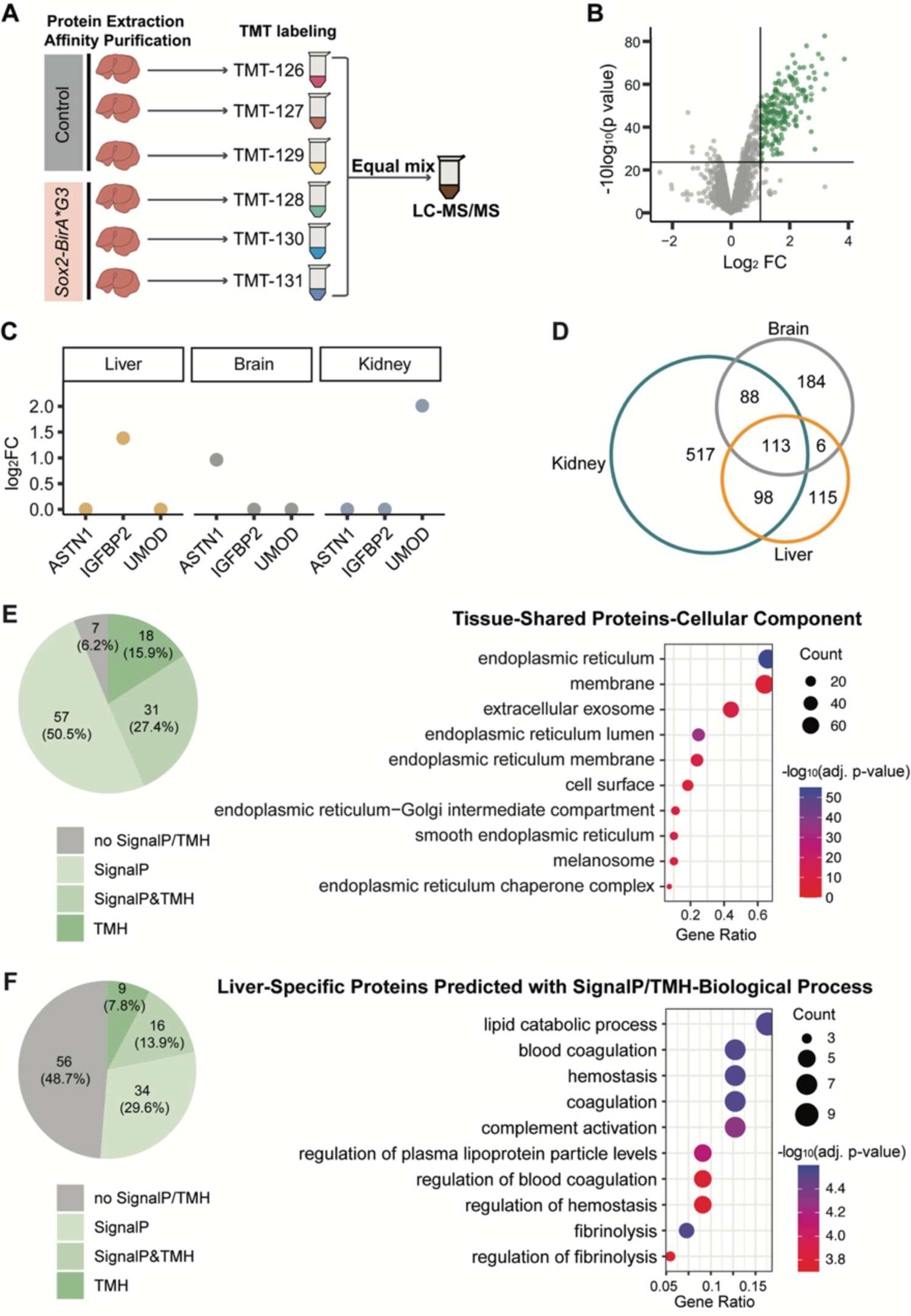
Identification of biotinylated proteins in *Sox2-BirA**\*G3*** liver, brain, and kidney tissues by mass spectrometry. (A) Representative schematic of TMT-based 6plex LC-MS/MS workflow for liver *Sox2-BirA*G3* (n=3) and control (n=3) samples. The same 6plex LC-MS/MS design was used for brain and kidney for individual MS runs. (B) Volcano plots of proteins detected in liver of *Sox2-BirA*G3* mice compared to control mice after streptavidin pulldown. Log_2_ FC were plotted on the x-axis and −10log_10_ (p value) were plotted on the y-axis. Significantly enriched proteins (adj. p-value< 0.05 and log_2_FC>1.0) in *Sox2-BirA*G3 A* mice compared to control mice are shown in green. (C) Relative abundance (log_2_FC) of representative proteins in liver, brain, and kidney. ES are labeled for the abundant proteins in their corresponding tissues. (D) Venn diagram showed the overlap of enriched proteins (ES method) among liver, brain, and kidney. (E) Shared enriched proteins among three tissues (113 proteins) were predicted with SignalP/TMH (left) and analyzed with DAVID analysis for cellular components annotation (right). Gene ratio indicates the percentage of genes annotated with the term over the total number of genes in the list. (F) Liver-specific enriched proteins (115 proteins) predicted with SignalP/TMH were analyzed with clusterProfiler (3.16.1) EnrichGO analysis. Left: a pie chart displayed the distribution of liver-specific proteins with SignalP/TMH prediction. Right: dot plots displayed the functional categorization of liver-specific enriched based on EnrichGO annotation, and the number of each category is displayed based on biological process. Gene ratio indicates the percentage of genes annotated with the term over the total number of genes in the list.

An enrichment score (ES) was generated as a measure of the abundance of proteins in *Sox2-BirA*G3* replicate samples compared to control samples, as previously described^13, 25^. Briefly, for 6plex TMT ratios of each tissue, TMT ratios were calculated by comparing each of the three *Sox2-BirA*G3* over each of the three control samples, giving rise to nine different datasets of TMT ratios. Then, we determined the false positive rate (FPR) based on proteins that are retained or traffic through the ER (positive control) and proteins that do not (negative control) (Supplementary Fig. 4D-E; see MS Hit Analysis in Methods). The number of TMT ratios for each protein that pass the TMT ratio cutoff based on an FPR of 0.1 was defined as enrichment score (ES) (Supplementary Fig. 4E). A protein where all its ratios pass the cutoffs has an enrichment score (ES) of 9 (highest confidence), where a protein where none of its ratios pass the cutoffs has a score of 0 (lowest confidence). Assigning an ES>=5 as the cutoff (Supplementary Fig. 4F) maximized the recovery of ER-targeted proteins with high specificity (Supplementary Fig. 4G-I).

In the conventional secretory pathway, proteins with either a signal peptide (SignalP) or transmembrane helix (TMH) travel through ER-Golgi apparatus. We used SignalP (v 5.0)^26–28^ and TMHMM (v 2.0)^29–31^ to predict the presence of SignalP and TMH on each tissue sample. Our analysis revealed that higher ES correlated with higher ratios of proteins with predicted SignalP/TMH (Extended Fig. 6D-F), suggesting that proteins with higher ES contain “hits” of higher confidence. Comparing across liver, brain, and kidney, 113 proteins were present in all three organ samples (Fig. 3D) and over 93% of these (106) showed a SignalP and/or TMH (Fig. 3E). Applying Gene Ontology (GO) functional category enrichment analysis to identify characteristic cellular component attributes of these tissue-shared proteins highlighted extracellular exosome and ER-Golgi terms in DAVID analysis (Fig. 3E; Extended Fig. 6G), consistent with ER localization of BirA*G3 and the expected labeling of proteins in the secretory pathway. These proteins are enriched in ER-Golgi related functions (Extended Fig. 6H).

TissueEnrich^32^, a tool for tissue-specific gene enrichment, was applied to liver, brain, and kidney-specific proteins (Extended Fig. 7A-C). Notably, all organ samples showed a strong enrichment profile for the expected organ type. Significant enrichment of SignalP/TMH was also seen in tissue-specific hits (Fig. 3F; Extended Fig. 7D, E). For biological process analysis, significantly enriched terms in liver showed unique features of liver function, which include lipid catabolic process and blood coagulation (Fig. 3F). Similarly, significantly enriched terms in brain showed unique features of brain function, such as synapse organization and assembly (Extended Fig. 7F). However, kidney-specific functional annotation terms were not observed amongst top terms in the kidney data (Extended Fig. 7G). The large majority of kidney enriched transporter and channel proteins are confined to short segments within the renal epithelium, organ-wide enrichment is likely to select for more secretory pathway proteins that are broadly distributed, or particularly abundant proteins such as UMOD, that are segmentally restricted (Fig. 3C; Extended Fig. 7E, G).

### Analysis of the serum secretomes of labeled tissues

To examine biotinylated proteins secreted into the serum of female *Sox2-BirA*G3* mice, serum from labeled mice was enriched using streptavidin-conjugated beads and bound fractions from secretomes of *Sox2-BirA*G3* (n=3) mice and associated controls (n=3) were analyzed by MS (6plex) (Fig. 4A; Extended Fig. 8A; Supplementary Data 4-5). PCA demonstrated tight clustering of the *Sox2-BirA*G3* group (Supplementary Fig. 5A-B). The affinity purified serum proteome identified by TMT-based MS has minimal background compared with tissue samples (Extended Fig. 8B). Accordingly, we used a log_2_ FC>1.0 and adj. p-value<0.05 to define *BirA*G3* serum enriched proteins over control serum to score enriched serum proteins from two independent MS runs of the same samples processed by two different research labs at different universities and run by the same MS group (plex1 and plex2).

**Fig. 4.**
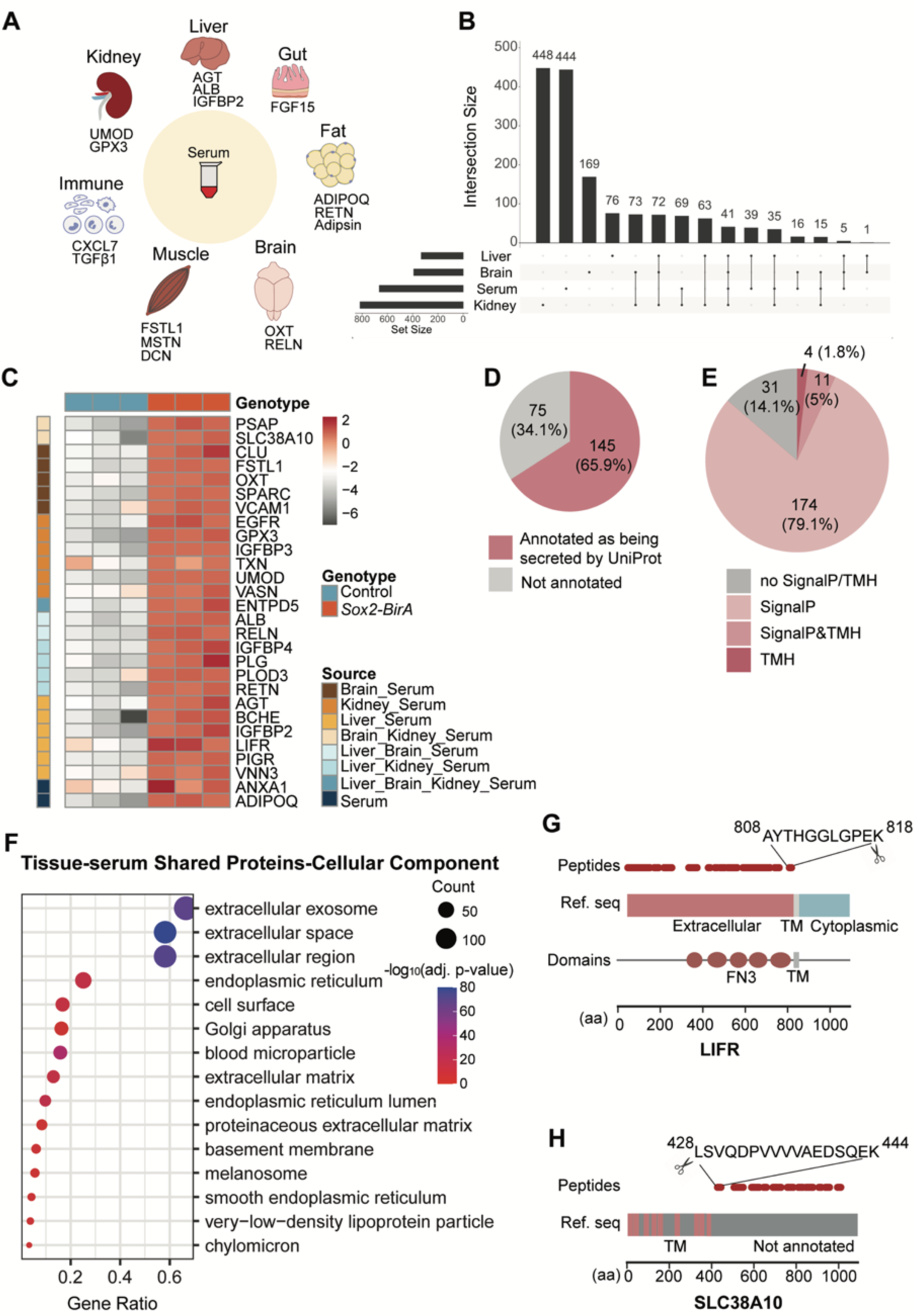
Analysis of biotinylated proteins secreted to peripheral blood. (A) Schematic of tissue secreted proteins identified by LC-MS/MS (6plex; *Sox2-BirA*G3* n=3, control n=3) from affinity purified serum from *Sox2-BirA*G3* mice. (B) Upset plot showed the overlap of enriched proteins among serum, liver, brain, and kidney. (C) Heatmap of representative proteins enriched in *Sox2-BirA*G3* serum relative to controls. Log_2_ expression values are shown by color and intensity of shading. Grey, low; red, high. (D) Pie chart displayed the number of shared enriched proteins between serum and three tissues (220 proteins) that are annotated as secreted by UniProt/NCBI. (E) Pie chart displayed the distribution of shared enriched proteins between serum and three tissues (220 proteins) with predicted SignalP/TMH. (F) Shared enriched proteins between serum and three tissues were analyzed with DAVID analysis for cellular component annotation. Gene ratio indicates the percentage of genes annotated with the term over the total number of genes in the list. (G) Schematic of detected peptides for LIFR mapped onto its respective reference sequences with SMART protein domain annotation. Reference sequence is annotated with extracellular, transmembrane (TM) and cytoplasmic based on UniProt topology information. Amino acid sequences of the most C-terminal peptide are labeled. FN3: fibronectin type 3. (H) Schematic of detected peptides for Slc38a10 mapped onto its respective reference sequences. Reference sequence is annotated with TM based on UniProt topology information. Amino acid sequences of the most N-terminal peptide are labeled.

The majority of serum proteins (70.6%) were identified in both MS runs (plexes) (Supplementary Fig. 5C), highlighting the reproducibility of this method. Most serum enriched proteins are annotated to contain a SignalP or TMH (64.8% of total and 76.6% of proteins shared between two plexes; Extended Fig. 8C) accounting for enriched cellular component terms for extracellular exosome/space/region (Supplementary Fig. 5D-E). As expected, biological process annotation for serum enriched proteins predicted with SignalP/TMH are enriched in immunity and complement-related terms (Supplementary Fig. 5F). To identify potential proteins secreted from tissues to serum, we compared the enriched proteins from serum with those from the three tissues and found 220 total overlapping proteins between serum and i) liver, ii) brain, or iii) kidney (Fig. 4B). A list of selective serum enriched proteins was shown in a heat map representing log_2_FC between *Sox2-BirA*G3* and control sample, including classic hepatocyte secreted proteins, such as Angiotensinogen (AGT) and Albumin (ALB), brain cell marker, neuron neuropeptide Oxytocin (OXT), as well as kidney markers, uromodulin (UMOD) and insulin-like growth factor-binding protein-3 (IGFBP3) (Fig. 4C).

Out of the 220 tissue-serum shared proteins, 65.9% were annotated as secreted by UniProt/NCBI (Fig. 4D), and the majority (85.9%) were predicted to have a SignalP or TMH (Fig. 4E). These tissue-serum shared proteins were then analyzed for GO term enrichment analysis (Fig. 4F; Extended Fig. 8D-E). DAVID analysis demonstrated significant enrichment in ‘extracellular’ annotations in the cellular component annotation (Fig. 4F). The biological process GO term for these tissue-shared proteins with predicted SignalP/TMH are enriched in coagulation and hemostasis (Extended Fig. 8E).

Ectodomain shedding is an important post-translational mechanism for regulating the function of cell surface proteins, which involves the proteolytic cleavage of transmembrane (TM) cell surface proteins, and release of circulating, soluble form^33, 34^. Five well-documented TM proteins (LIFR, EGFR, VCAM1, Slc38a10 and PIGR) appeared in our serum proteomic datasets. For each, sequenced peptides exclusively mapped to the annotated extracellular domains, consistent with ectodomain shedding (Fig. 4G; Extended Fig. 8G-I). While ectodomain shedding has been reported for LIF receptor subunit α (LIFR)^35^, polymeric immunoglobulin receptor (PIGR)^36^, epidermal growth factor receptor (EGFR)^37^, and vascular cell adhesion molecule 1 (VCAM1)^38^, solute carrier family 38 member 10 (Slc38a10) have not been directly linked to shedding, though an extramembrane fragment of human Slc38a10 has been reported in plasma^39^ (Fig. 4G-H; Extended Fig. 8F-H). Approximate protein cleavage sites can be assigned on the basis of the most N- or C-terminal peptides (Fig. 4G-H; Extended Fig. 8F-H). For example, the in vivo cleavage site for LIFR is predicted to be between the fifth fibronectin type 3 (FN3) domain and the TM domain, similar to a previous report^15^. Intriguingly, we also detected thioredoxin (TXN) and Annexin1 (ANXA1) in our dataset (Fig. 4C), each of which are reported to be released from the cell through unconventional protein secretion (UPS)^40–42^, suggesting this method can give further insight into UPS, with potentially other candidates among the detected proteins lacking a predicted SignalP or TMH domain.

To confirm key hits in these secreted serum protein data, we performed western blot analysis of 11 proteins observed in the MS analyses of biotinylated serum proteins (Extended Fig. 8I, J). These validation analyses were carried out in both male and female mice (Extended Fig. 8J). Consistent with the MS data, all 11 proteins were enriched in *Sox2-BirA*G3* serum after affinity purification.

### Biotin labeling of the hepatocyte secretome

We next applied and extended the approach to determine the feasibility of identifying a tissue cell type specific secretome. For this, we utilized *Alb-Cre* to enable hepatocyte-specific biotinylation of proteins in the liver^62^. Biotin chow was given to adult *Alb-Cre; BirA*G3* (abbreviated as *Alb-BirA*G3*) and control mice for 5 days, and then well-perfused tissues were collected after exsanguination (Supplementary Fig. 6A). mKate2 was specifically detected in the *Alb-BirA*G3* liver (Fig. 5A; Extended Fig. 9A; Supplementary Fig. 6B). BirA*G3 showed a homogenous distribution restricted to the hepatocytes of *Alb-BirA*G3* liver (Fig. 5B). BirA*G3 activity in *Alb-BirA*G3* liver enabled specific protein biotinylation in liver protein lysates to be compared with control mice (Fig. 5C). *Alb-BirA*G3* serum samples showed strong, specific protein biotinylation relative to serum from biotin administered controls lacking active BirA*G3 (Fig. 5D). The biotinylation banding patterns of affinity purified biotinylated proteins were comparable to those observed in total protein lysates of liver and serum samples (Fig. 5C-D; Extended Fig. 9B-F). Silver staining of affinity purified biotinylated proteins showed specific bands in *Alb-BirA*G3* serum, with similar size distributions between sexes (Fig. 5E).

**Fig. 5.**
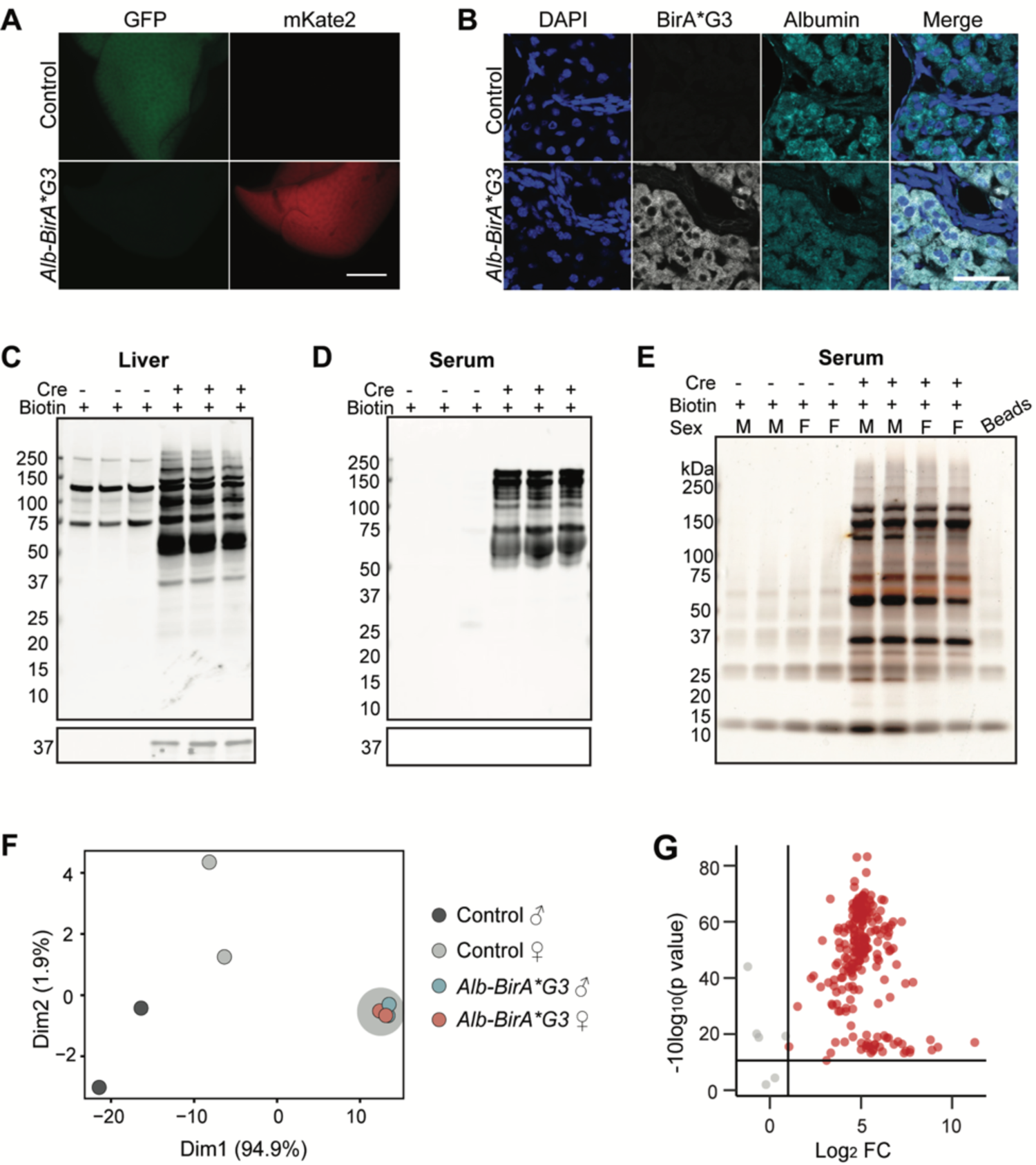
Generation and characterization of *Alb-BirA**\*G3*** mice. (A) Whole mount images of native GFP and mKate2 fluorescence in livers of *Alb-BirA*G3* and control mice. (Scale Bar: 3mm) (B) Immunofluorescence staining shows expression of BirA*G3 and hepatocyte marker Albumin in the liver sections from *Alb-BirA*G3* and control mice. Scale Bar: 50μm. (C-D) Western blotting of total protein from liver (C) and serum (D) in *Alb-BirA*G3* mice compared to control mice. Upper: Streptavidin labeling. Lower: BirA*G3 (∼35kDa). Each lane is a biological replicate from individual mice (n=3/genotype). (E) Silver stain of affinity purified biotinylated proteins from serum in *Alb-BirA*G3* mice compared to control mice. Each lane is a biological replicate from individual mice (n=4/genotype). (F) PCA of streptavidin-purified serum proteins from *Alb-BirA*G3* and control mice. Each dot represents a sample, which is colored by the annotation of mouse genotype and sex. *Alb-BirA*G3* samples are manually circled by grey shadow. (G) Volcano plots of proteins detected in serum of *Alb-BirA*G3* mice compared to control mice after streptavidin pulldown in mass spectrometry. Log_2_ FC were plotted on the x-axis and −10log_10_ (p value) were plotted on the y-axis. Significantly enriched proteins (adj. p-value< 0.05 and log_2_FC>1) in *Alb-BirA*G3* mice compared to control mice are shown in red.

### Identification of hepatocyte secretome

Streptavidin purified serum proteins of *Alb-BirA*G3* (n=4) and control (n=4) mice were analyzed by LC-MS/MS (8plex) to identify secreted hepatocyte proteins (Supplementary Data 6). PCA demonstrated male (n=2) and female (n=2) *Alb-BirA*G3* samples cluster together, indicating similar hepatocyte secretome between the two sexes (Fig. 5F). Approximately 80% of proteins (181/189) were specifically enriched (log_2_FC>1.0 and adj. p-value<0.05) in the *Alb-BirA*G3* group (Fig. 5G). A heatmap showed very low protein abundance in the control group, consistent with the absence of biotinylated proteins in the control serum samples (Fig. 6A; Fig. 5E). To determine cell labeling specificity, TissueEnrich on serum enriched proteins in the *Alb-BirA*G3* group showed a highly liver-specific organ profile using the mouse ENCODE dataset (Fig. 6B).

**Figure 6.**
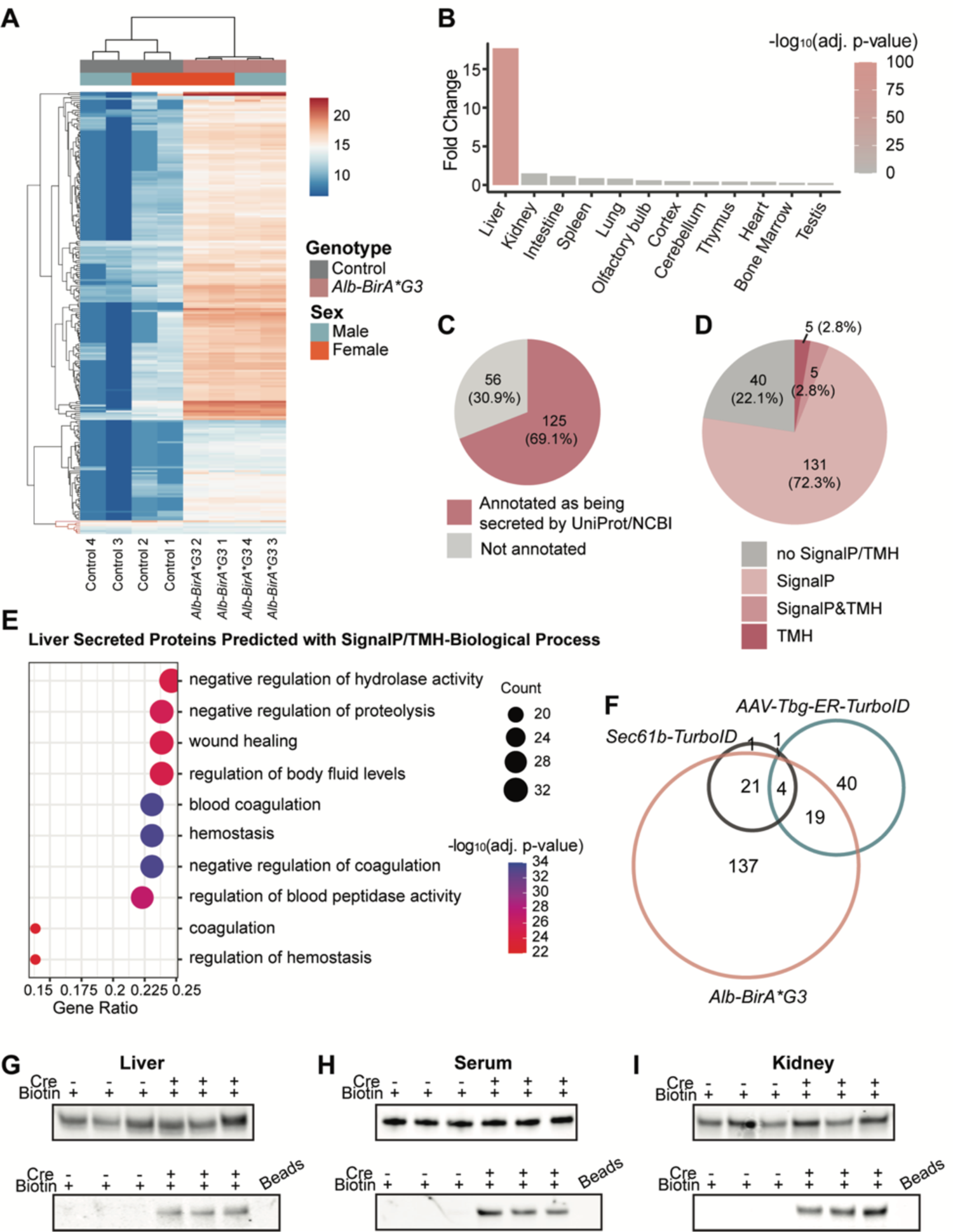
Analysis of hepatocyte-secreted serum proteins in *Alb-BirA*G3* mice. (A) Cluster heatmap of proteins enriched in *Alb-BirA*G3* serum relative to controls. Log_2_ expression values are shown by color and intensity of shading. Blue, low; red, high. Serum enriched proteins in the *Alb-BirA*G3* group are highlighted in black color in the hierarchical clustering. (B) Enriched proteins in *Alb-BirA*G3* serum were analyzed with TissueEnrich to calculate tissue-specific gene enrichment. (C) Pie chart displayed the number of enriched proteins in *Alb-BirA*G3* serum that are annotated as secreted by UniProt/NCBI. (D) Pie chart displayed the distribution of enriched proteins in *Alb-BirA*G3* serum with predicted SignalP/TMH. (E) Enriched proteins in *Alb-BirA*G3* serum predicted with SignalP/TMH were analyzed with clusterProfiler (3.16.1) EnrichGO analysis for biological process. Gene ratio indicates the percentage of genes annotated with the term over the total number of genes in the list. (F) Area-proportional venn diagram showed the overlap among enriched proteins in *Alb-BirA*G3* serum, *AAV-Tgb-ER-TurboID* plasma^15^, and *Sec61-TurboID* plasma^14^. (G-I) AGT levels in total protein lysates (upper) and affinity purified biotinylated proteins (lower) from liver (F), serum (G), and kidney (H) in *Alb-BirA*G3* mice compared to control mice. Input: 500μg proteins. Bead lanes are affinity purification negative control without protein input. Each lane is a biological replicate from individual mice (n=3/genotype).

Over half (69.1%) of the proteins enriched in *Alb-BirA*G3* serum were annotated as secreted and a majority (77.9%) of these proteins were predicted with SignalP/TMH (Fig. 6C, D). Functional annotation showed significant enrichment for ‘extracellular’ terms (Extended Fig. 10A-B). Biological process annotation for the subset of serum enriched proteins with predicted SignalP or TMH, specific to the *Alb-BirA*G3* group demonstrated enriched terms for key liver protein functions: coagulation and hemostasis (Fig. 6E)^41^.

Recently, other groups have used adeno-associated virus carrying *ER-TurboID* under the control of the hepatocyte-selective *Tbg* (thyroxine binding protein) promoter (*AAV-Tbg-ER-TurboID*)^15^ and adenovirus carrying *Sec61b* (Protein transport protein Sec61 subunit beta)*-TurboID*^14^ to label the liver secretome of mice. While these two previous datasets only share 5 proteins, both displayed a significant overlap (23/64 and 25/27, respectively) with our *Alb-BirA*G3* dataset, highlighting the power of the transgenic strain (Fig. 6F). Further, the 137 proteins uniquely detected in the *Alb-BirA*G3* dataset displayed a strong correlation with liver-specific proteins and the majority (67.2%) was as predicted with SignalP/TMH (Extended Fig. 10C, D). Well-characterized hepatocyte secreted proteins, including APOE, AGT, IGFBP2, were only seen in the *Alb-BirA*G3* unique dataset, highlighting an improved coverage of secreted liver proteins using this strategy.

Finally, we asked if secreted proteins can be labeled in the organ of origin and subsequently detected in a destination organ after affinity purification. The renin-angiotensin-aldosterone system regulates blood pressure in conjunction with the liver, kidney, and lung^43^. In this pathway, angiotensinogen (AGT) is produced by the liver and sequentially converted by kidney and lung enzymes to vasoconstrictive angiotensin II (AGTII). Liver released AGT is reabsorbed from the renal filtrate by proximal tubule cells in the kidney^44^. Consistent with this view, AGT was detected in the liver, serum, and kidney total protein from both *Alb-BirA*G3* and control mice (Fig. 6G-I), but following streptavidin affinity purification, AGT was only detected in the liver, serum, and kidney from *Alb-BirA*G3* mice, but not in control mice (Fig. 6G-I). Together, these data support the trafficking of AGT produced in liver through the blood stream with uptake in the kidney.

## Discussion

The genetic system presented here allows for rapid (within 1 hour) and broad (representative cell types) *in vivo* biotinylation of proteins within the secretory pathway of the living mouse. Long term studies of BirA*G3 production show no obvious pathology. Although we do observe cell-type variability in BirA*G3 levels, that may be countered using tissue or time-dependent CRE activator lines as we show here comparing *Sox2-BirA*G3* with *CAGG-BirA*G3*.

TMT-based LC-MS/MS captured a number of well-characterized secreted proteins in *Sox2-BirA*G3* serum, including proteins with hormone like properties, known to circulate at µg/mL (ADIPOQ^45^) and ng/mL (ANGPTL^46^, MST1^47^, MSTN^48^, RETN^45^, CXCL7^49^, IGF1^50^, FGF15^51^) levels. Interestingly, these represent examples of proteins derived from multiple tissues: adipose tissue (ADIPOQ, Adipsin^52^, and RETN), muscle (MSTN), immune cells (CXCL7), liver (IGF1), and intestine (FGF15). Furthermore, the approach provided insight into ectodomain shedding and UPS.

Three research groups have reported various applications of proximity ligation techniques to profile the mammalian secretome. Two groups^14, 15^ profiled the mouse hepatocyte secretome via viral delivery of TurboID. Viral vectors however present unavoidable problems, such as viral tropism, which limits the use in some tissues. For instance, kidney has been quite difficult to transduce with any viral vector currently available^53^. For brain, viruses need to be intracranially administered due to the blood-brain barrier, for higher transduction efficiency and brain-specific targeting and viral vectors have broad tropisms for neural and glial cell types^18, 19^. Directly comparing the *Alb-BirA*G3* mouse strain to the two viral studies profiling the hepatocyte secretome, our study showed an improved coverage for relevant hepatocyte proteins which may reflect the stable and efficient expression of BirA*G3, as well as other differences in the downstream analytical pipeline.

A third group^17^ reported a transgenic mouse expressing BioID, a less efficient biotin ligase (as shown in various studies^12, 17, 54^), in the endoplasmic reticulum. Characterization of the endothelial and muscle secretome showed few secreted proteins, likely due to the low activity of BioID2 in ER lumen^12^. Furthermore, the kinetics of labeling are reported to be much faster with BirA*G3, suggesting the line reported here will be better suited to dynamic labeling studies^12^.

Incomplete retention of TurboID-KDEL in the endoplasmic reticulum has been documented^14^. In our MS data, we identified tryptic peptides for BirA*G3 in both *Sox2-BirA*G3* and *Alb-BirA*G3* serum. However, using western blotting, we did not detect BirA*G3 in total serum proteins or affinity purified proteins in *Sox2-BirA*G3* serum (Fig. 2C; Extended Fig. 5D). Biotinylation, with biotin ligase, is an ATP-dependent reaction, and ATP is normally restricted within the cell^55^. Biotin concentration within the cell and the concentration of proteins in the secretory pathway, likely also impact the efficiency of the enzymatic reaction. Together, these kinetic arguments make it unlikely there is a significant impact from low levels of circulating BirA*G3 on biotinylation of serum proteins. This is supported by the *Alb-BirA*G3* data which demonstrated a robust biotin profile for specific liver-derived proteins with no labeling of abundant proteins secreted by other organs, identified in the *Sox2-BirA*G3* study.

Recent studies have found that many cytosolic proteins lacking a SignalP/TMH (leaderless cargoes) are released through a type III UPS mechanism into the ER^40^. ANXA1 is one such protein detected in our serum dataset (Fig. 4C)^40^ suggesting further exploration of potential unconventional secretion processes for other leaderless proteins in our data. Recent studies show that approximately 13% of the human protein-coding genes encode for roughly 2640 secreted proteins, but relatively few have been functionally annotated and characterized^4^. Since many protein functions are conserved within mammals, we anticipate our platform will serve as a valuable resource for deciphering the mammalian secretome, under healthy or diseased conditions.

## Acknowledgements

This work was supported by a NIH Transformative R01 grant 5R01DK121409 to A.P.M., A.Y.T., S.A.C., and N.P. We thank the Janelia Gene Targeting and Transgenics Facility (Director: Caiying Guo) for the BirA*G3 mouse generation. We thank Peter Calabrese and Ivy Xiong from University of Southern California (USC), Pierre Michel Jean Beltran from Broad Institute, Xianwen Ren from Peking University, Kevin Blighe from Rancho BioSciences, and Geetu Tuteja from Iowa State University for help with mass spectrometry data analysis. We thank Jeffrey Boyd and Bernadette Masinsin from the Flow Cytometry Facility of Eli and Edythe Broad CIRM Center for Regenerative Medicine and Stem Cell Research at USC for assisting with hepatocyte sorting. We thank Seth Ruffins from Optical Imaging Facility and Riana Parvez of Eli and Edythe Broad CIRM Center for Regenerative Medicine and Stem Cell Research at USC for assisting with fluorescence imaging. We thank Genome Technology Access Center from Washington University for performing bulk RNA-seq. Norbert Perrimon is an investigator of the Howard Hughes Medical Institute.

## Contributions

A.P.M., R.Y., A.S.M., I.A.D., N.P., S.A.C., N.D.U., A.T., and J.W. designed the experiments. A.P.M., R.Y., A.S.M., and I.A.D. wrote the manuscript with input from all authors. R.Y., A.S.M., J.G., J.M., and D.R. performed the experiments. Mass spectrometry-based proteomic analyses were carried out at the Broad Institute by N.D.U., D.K.C., and C.X., and at UCLA by Y.H. and J.S. R.Y., A.S.M., I.A.D., Y.H., N.D.U., D.K.C., C.X., F.Q., J.S., and S.Q. analyzed proteomics data. A.S.M. analyzed bulk RNA-seq data.

## Materials and Method

### Animal studies

Institutional Animal Care and Use Committees (IACUC) at the University of Southern California reviewed and approved all animal work as performed in this study. All work adhered to institutional guidelines. mESCs (B6(Cg)-Tyr/J, Stock No.: 000058, The Jackson Laboratory)^56^ carrying the loxP-flanked *BirA*G3* were aggregated with 8-cell C57bl/6J embryos to obtain chimeras. The resulting chimeric mice were bred with R26PhiC31 (B6.129S4-*Gt(ROSA)26Sor^tm3(phiC31*)Sor^*/J, Stock No.: 007743, The Jackson Laboratory, backcrossed to C57bl/6J for 13 generations)^57^ females to remove attB-neoR-attP cassette. The resulting mice carry the *BirA*G3* allele, namely *BirA*G3* mice. The *Sox2-Cre* mice (B6.Cg-*Edil3^Tg(Sox2-cre)1Amc^*/J, stock no.: 008454, The Jackson Laboratory)^23^, the *TdTomato* mice (B6.Cg-*Gt(ROSA)^26Sortm14(CAG-tdTomato)Hze^*/J, Stock No.: 007914, The Jackson Laboratory)^24^, the *CAGGCre-ER^TM^* mice(FVB.Cg-*^Tg(CAG-cre/Esr1*)5Amc^*/J, stock no.: 017595, The Jackson Laboratory) ^58^, and the *Alb-Cre* mice (B6.Cg-*Speer6-ps1^Tg(Alb-cre)21Mgn^*/J, stock no.: 003574)^59^ were used as described previously.

### In vivo Assays

For all *in vivo* assays, tissues were collected as follows. Mice were euthanized at 8-32 weeks, blood was then collected from the inferior vena cava, followed by perfusion with 1×cold DPBS (Dulbecco’s Phosphate Buffered Saline). The blood was allowed to clot at room temperature for 30 minutes and then spun down at 2,000 x g for 15 minutes at 4°C. The serum was collected and spun again at 2,000 x g for 15 minutes at 4°C, then removed to a fresh tube and flash frozen (in liquid nitrogen) before being stored at −80°C until being used. After perfusion, tissues were collected and rinsed in 1×cold DPBS before being minced with a razor blade and aliquoted into tubes. Tissues were then flash frozen and stored at −80°C until being used.

For the biotin administration method study, *Sox2-BirA*G3* and control mice at 8-12 weeks were given biotin via water (n=2) (5mM, pH 7.4; Sigma B4639-5G), chow (n=2) (2,000ppm; LabDiet, SWLP), or both (n=2) for 7 days. After 7 days, serum was collected and stored as described above.

For the biotin and Cre dependence labeling study, *Sox2-BirA*G3* and control mice at 8-12 weeks were given biotin chow (2,000ppm) or normal diet (LabDiet 5053) for 7 days. After 7 days, serum, liver, brain, and kidney were collected as described above.

For the BirA*G3 temporal labeling assay, *Sox2-BirA*G3* (n=3/timepoint) and control mice (n=3) at 8-12 weeks were given biotin by subcutaneous injection (100μL 180mM sterile biotin water, pH 7.4) and water (5mM, pH 7.4) after the injection until collection. Serum, liver, kidney, and brain were collected from mice at 1, 2, 4, 6, 10, and 12 hours post-injection. Serum was collected as described above.

For all other mouse studies, *Sox2-BirA*G3* and control mice at 8-32 weeks were given biotin chow (2,000ppm) for 5 days. Tissues were then collected as described above.

For ERT2 mice, tamoxifen (2mg/40g body weight in corn oil; Sigma cat. T5748-1G) was administered by abdominal injection twice, 3 days apart prior to biotin studies.

### In vitro Assays

To generate endoplasmic reticulum stress positive controls 80% confluent mouse 3T3 cells (NIH/3T3 ATCC cat. CRL-1658) were treated with 5ug/mL Tunicamycin (R&Dsystems 590507) in DMSO for 5 hours. Cells were then briefly rinsed with 1X DPBS and then scrapped off the plate in fresh DPBS. Cells were pelleted by centrifugation at 300 x g for 10 minutes. Supernatant was then removed, and dry cell pellets were snap frozen on dry ice and then stored at −80C until used. For high-resolution calnexin-BirA staining, mouse embryonic fibroblasts (MEFs) isolated from *Sox2-BirA*G3* (n=1) and control (*Sox2-Cre)* (n=1) mice were cultured on coverslips coated with 0.1% gelatin. Cells at 70% confluence were briefly rinsed in 1X DPBS and then fixed in 4% PFA for 10 minutes followed by 3 washes in 1X DPBS. Coverslips with cells were stored at 4°C in 1X DPBS in the dark until used.

### Hematoxylin and Eosin staining

Tissues (n=3/genotype) were collected and fixed in 10% phosphate-buffered formalin overnight at 4°C. Paraffin sections were prepared using standard procedures. The tissue sections were deparaffinized by immersing in xylene and rehydrated through graded alcohol series, dyed with hematoxylin and eosin (H&E) and then rinsed with water. All slices were dyed with hematoxylin and eosin (H&E) and then rinsed with water. Each slide was dehydrated through graded alcohols. Tissue sections were finally soaked in xylene twice. The sections were examined under a light microscope for evaluation of pathological changes.

### Frozen Tissue Preparation and Sectioning

Briefly, tissues were harvested from DPBS-perfused mice and then fixed in 4% paraformaldehyde for 2 hours at 4°C. Tissues (except brain: fixation overnight and 30% sucrose for 48 hours) were then washed 3 times in 1X DPBS with calcium and magnesium before being incubated in 30% sucrose overnight. The following day, tissues were washed in OCT 3 times to remove excess sucrose and then embedded in OCT (VWR, 25608-930) and frozen in a dry-ice ethanol bath before being stored at −80°C. Tissue blocks were thawed to −20°C overnight and then cryosectioned at 10-16μm at −20°C and placed on glass slides. Slides were then stored at −80°C until immunostained.

### Immunofluorescent Staining and Confocal Microscopy

Frozen sectioned tissues were thawed at room temp for 10 minutes. Slides were then rinsed in 1× DPBS with calcium and magnesium for 10 minutes. Coverslips with cells were removed from 4°C 1X DPBS and treated as slides. Slides were permeabilized in 0.25% Triton-X100 (Sigma X100-500ML) for 5 minutes, then incubated in blocking buffer (2.0% Sea Block (Thermo, 37527) + 0.125% Triton-X100 in 1× DPBS with calcium and magnesium) for 1 hour at room temperature. Slides were then incubated in primary antibody (Supplementary Table 1) diluted in blocking buffer, overnight at 4°C. The following day, primary antibodies were removed, and slides were washed in blocking buffer four times for 5 minutes each. Slides were then incubated in secondary antibody diluted in blocking buffer for 1 hour at room temperature. Secondary antibody (Supplementary Table 2) was removed, and slides were washed in blocking buffer four times for 5 minutes each. Slides were then incubated in 1 mg/mL Hoechst 33342 (Thermo, H3570) in 1× DPBS with calcium and magnesium for 10 minutes at room temperature. Slides were then washed twice in 1× DPBS with calcium and magnesium for 5 minutes each. Slides were mounted in Immu-Mount (Thermo, 9990402) and imaged at 40× or 63× using the Leica SP8 confocal microscope. Brain images are median intensity projections from z-stacks of 0.5-1.0μm steps except the whole brain tiles scans. All images presented represent at least three images per tissue and 2 biological samples per genotype. Whole tissue section scans were imaged using the Zeiss AxioScan Z1 Slide Scanner at 20× to generate high-resolution tiled images of tissues sections.

### Hepatocyte isolation and qPCR

We used the classic two-step collagenase perfusion technique to isolate primary mouse hepatocytes. Briefly, The *Sox2-BirA*G3* and control livers were perfused by perfusion medium (GIBCO #17701-038) through Inferior vena cava. Then, the livers were dissociated by liver digested medium (GIBCO 17703-034), which is incubated 30-60 min in 37°C before use. mKate2^low^ and mKate^high^ hepatocytes were sorted by flow cytometry based on mKate2 levels. Total RNA was extracted using RNeasy mini kit (Qiagen, 74104). cDNA was synthesized using SuperScript™ IV VILO™ Master Mix (Thermo #11766050). qPCR was performed with Luna Universal qPCR Master Mix Protocol (New England Biolab #M3003) on a Roche LightCycler 96 System. Primers used in RT-qPCR are listed as follows: *BirA*G3:*

Forward: CTCCCCGTGGTTGACTCTAC Reverse: CTCCCCGTGGTTGACTCTAC

Alb:

Forward: GTCTTAGTGAGGTGGAGCATGACAC Reverse: GCAAGTCTCAGCAACAGGGATACAG

Gapdh:

Forward: CATGGCCTTCCGTGTTCCTA Reverse: CCTGCTTCACCACCTTCTTGAT

The delta-delta Ct method, also known as the 2^-ΔΔCt^ method, was used to calculate the relative fold gene expression.

### RNA Sequencing and Analysis

Mouse samples were collected for RNA as described above. After collecting samples were immediately stored in RNALater (Sigma 90901-100ML) at 4°C overnight before being stored at −80°C until processing. Before RNA extraction samples were removed from RNALater and briefly rinsed in RNase free dH_2_O. Total RNA was then prepared from *Sox2-BirA*G3* and control liver, kidney, and whole brain (n=2/genotype, all female) using RNeasy mini kit (Qiagen 74104) according to kit instructions with the following modifications. For whole brain, brains were homogenized in 1400μL RLT buffer. Brain samples were then further diluted 1:25 in fresh RLT buffer (total final volume 350μL) to avoid overloading the columns. Liver and kidney tissue samples were processed exactly following kit instructions. RNA integrity was determined using Agilent TapeStation 4200. Samples were then sequenced by Washington University in St. Louis School of Medicine Genome Technology Access Center (GTAC) using the following methods. Total RNA integrity was determined using Agilent BioAnalyzer. Library preparation was performed with 10ng of total RNA wth Bioanalyzer RIN score of greater than 8.0. ds-cDNA was prepared using the SMARTer Ultra Low RNA kit from Illumina Sequencing (Takar-Clonetech) per manufacturer’s protocol. cDNA was fragmented using a Covaris E220 sonicator using peak incident power 18, duty factor 20%, cycles per burst 50 for 120 seconds. cDNA was blunt ended, had A base added to the 3’ ends, and then had Illumina sequencing adapters ligated to the ends. Ligated fragments were then amplified for 12-15 cycles using primers incorporating unique dual index tags. Fragments were sequenced on an Illumina NovaSeq-6000 using paired end reads extending 150 bases. Raw paired end read files were first trimmed using Trimmomatic v0.38 and then aligned to GRCm39 using STAR v2.7.0e with default options. Read counts were quantified using RSubread_2.2.6 (featureCounts) without multimapping. Read counts were then analyzed using DESeq2 v1.28.1 in R (v4.0.0) using standard approaches with cutoffs of log_2_ fold change > 2.0 and adjusted p-value < 0.05.

### Protein lysate preparation

Protein lysates were prepared as described previously^13^ with the following modifications. homogenized in 500μL RIPA complete lysis buffer (RIPA buffer (ThermoFisher, 89901) with 1× cOmplete EDTA-free protease inhibitor cocktail (Sigma, 11873580001), 1mM benzamidine hydrochloride (VWR, TCB0013-100G), 4μM pepstatin A (Sigma, EI10), 100μM PMSF (Sigma, 11359061001)) and bead homogenized using stainless steel beads (NextAdvance, SSB14B-RNA) for 5 minutes at setting 10, Bullet Blender Storm (NextAdvance, BT24M). Samples were then centrifuged at 14,000×g for 15 minutes at 4°C. Supernatants were transferred to protein loBind (Eppendorf) tubes. Protein lysate concentrations were determined using Pierce BCA (Thermo cat. 23227) microplate assay per manufacturer’s instructions. Lysates were then stored at −80°C.

### Streptavidin beads pulldowns

Streptavidin pulldowns were performed as described previously^13^ with modifications. Streptavidin magnetic beads (Thermo cat. 88817) were resuspended in lysis buffer (above) by magnetic separation (BioRad 1614916). We tested a series of volumes of beads and washing conditions and found that 5μL beads per 100μg protein together with the following washing conditions are sufficient (results available upon request). Pulldown reactions were set up in 450μL lysis buffer with 5μL beads per 100μg protein. Pulldowns were then incubated overnight at 4°C in a wheel rotator. The following day, pulldown reactions were washed 2× in lysis buffer, then 1× in 2M Urea in 10mM Tris, and finally 2× lysis buffer. After the final wash, lysis buffer was removed and beads were either boiled in 12μL 1× loading buffer (Li-Cor 928-40004) with 1.43 M β-mercaptoethanol or resuspended in 100μL lysis buffer and flash frozen, for western blotting and mass spectrometry respectfully.

### Fluorescent Western Blot Analysis

Western blots were performed with standard protocols and the following modifications. Equal amounts of total protein lysate were loaded per sample per reaction with 1× Li-Cor loading buffer (Li-Cor, 928-40004) with 1.43 M β-mercaptoethanol. For streptavidin pulldowns, beads were resuspended in 12μL 1× Li-Cor loading buffer (Li-Cor, 928-40004) with 1.43 M β-mercaptoethanol. All samples were then boiled at 95°C for 5 minutes to elute, then briefly spun down and kept on ice prior to loading. Total protein samples and pulldown elutes were loaded on 10% SDS acrylamide gels and ran in standard 1× SDS-Running buffer with Li-Cor 5μL one-color molecular marker (Li-Cor, 928-40000) at 60V for 30 minutes, followed by 120V for ∼50 minutes or until loading dye ran off. Samples on the gel were transferred to methanol activated PVDF 0.45μm membranes using BioRad’s wet tank mini-protean system for 1-3 hours at 250-300 constant mA in a sample dependent context. After transfer, membranes were dried at 37°C for 5 minutes and then re-activated with methanol. Blots were stained with Li-Cor’s Revert-700 Total Protein Stain (Li-Cor, 926-11010) for normalization and imaged using a Li-Cor Odyssey Clx. Blots were then de-stained per kit instructions and put in block (Li-Cor Intercept block, 927-60001) for 1 hour, room temperature, shaking. Blots were then transferred to primary antibody (Supplementary Table 1) (block with 0.2% Tween20) overnight at 4°C, shaking. The following day, blots were washed four times in TBS-T for 5 minutes each at room temperature, shaking, and then incubated in secondary antibody (Supplementary Table 2) in block with 0.2% Tween20 and 0.1% SDS, and/or streptavidin conjugate (1:5,000; 680 or 800, Li-Cor, 926-68079, 926-32230) if visualizing biotinylated proteins, for 1 hour at room temperature, shaking. Blots were then washed twice with TBS-T for 5 minutes each, room temperature, shaking, followed by two 5-minute TBS washes at room temperature, shaking. Blots were imaged on a Li-Cor Odyssey Clx using Li-Cor’s ImageStudio (Version 5.2.5). After imaging blots were dried at 37°C for 5 minutes, then stored. For phosphorylated proteins (EIF2α), proteins were blotted for the phosphorylated state as described above. After imaging, phospho-blots were stripped is 10mLs 1X Restore Fluorescent Western Blot stripping buffer (Thermo cat. 62300) for 20 minutes at room temperature, shaking. Blots were then briefly rinsed with dH_2_O twice, re-blocked for 30 minutes at room temperature shaking, before primary incubation with the total protein antibody and secondary as described above. All western blot images were exported from Li-Cor, pseudo-colored and converted to RGB tiffs in ImageJ (v1.51S) for figures. For specific proteins, bands were selected based on molecular weight from antibody manufacturer information and literature. Note that all western blot experiments were repeated at least twice with different biological samples and produced consistent results.

### Fluorescent Western Blot Quantification

Biotinylation levels and proteins of interest were quantified via western blot using Li-Cor’s fluorescent western blot ImageStudio (Version 5.2.5) and Emperia Studio (Version 1.3.0.83) analysis software and protocols. Total protein stain images of each blot were used to normalize biotinylation (streptavidin) or protein of interest signal intensity in RStudio (Version 1.3.959, R Version 4.0.0) by determining the lane normalization factor (Li-Cor protocol) for each blot per manufacturer’s instructions. ggplot2 (Version 3. 3.5) and GraphPad Prism 9.0 were used to visualize normalized biotinylated protein signal.

### *Sox2-BirA\**G3 Analysis by MS (corresponding to **Figure 3-4** and Extended Figures 6-8)

After streptavidin beads pulldowns, *Sox2-BirA*G3* and control samples were sent to The Broad Institute of Harvard and MIT for MS.

#### i. On-bead digestion

Samples collected and enriched with streptavidin magnetic beads were washed twice with 200 μL of 50mM Tris-HCl buffer (pH 7.5), transferred into new 1.5 mL Eppendorf tubes, and washed 2 more times with 200 μL of 50mM Tris (pH 7.5) buffer. Samples were incubated in 0.4 μg trypsin in 80 μL of 2M urea/50mM Tris buffer with 1 mM DTT, for 1 h at room temperature while shaking at 1000 rpm. Following pre-digestion, 80 μL of each supernatant was transferred into new tubes. Beads were then incubated in 80 uL of the same digestion buffer for 30 min while shaking at 1000rpm. Supernatant was transferred to the tube containing the previous elution. Beads were washed twice with 60 μL of 2M urea/50mM Tris buffer, and these washes were combined with the supernatant. The eluates were spun down at 5000 × g for 1 min and the supernatant was transferred to a new tube. Samples were reduced with 4 mM DTT for 30 min at room temperature, with shaking. Following reduction, samples were alkylated with 10mM iodoacetamide for 45 min in the dark at room temperature. An additional 0.5 μg of trypsin was added and samples were digested overnight at room temperature while shaking at 700 × g. Following overnight digestion, samples were acidified (pH < 3) with neat formic acid (FA), to a final concentration of 1% FA. Samples were spun down and desalted on C18 StageTips as previously described56. Eluted peptides were dried to completion and stored at −80 °C.

#### ii. TMT labeling of peptides

Desalted peptides were labeled with TMT (6-plex) reagents (ThermoFisher Scientific). Peptides were resuspended in 80 μL of 50 mM HEPES and labeled with 20 uL 20mg/mL TMT6 reagents in ACN. Samples were incubated at RT for 1 h with shaking at 1000 × rpm. TMT reaction was quenched with 4 μL of 5% hydroxylamine at room temperature for 15min with shaking. TMT labeled samples were combined, dried to completion, reconstituted in 100 μL of 0.1% FA, and desalted on StageTips.

#### iii. bRP stage tip fractionation of peptides

50% of the TMT labeled peptide sample was fractionated by basic reverse phase (bRP) fractionation. StageTips packed with 3 disks of SDB-RPS (Empore) material. StageTips were conditioned with 100 μL of 100% MeOH, followed by 100 μL 50% MeCN/0.1% FA and two washes with 100 μL 0.1% FA. Peptide samples were resuspended in 200 μL 1% FA (pH<3) and loaded onto StageTips. 6 step-wise elutions were carried out in 100 μL 20 mM ammonium formate buffer with increasing concentration of 5%, 10%, 15%, 20%, 25%, and 45% MeCN. Eluted fractions were dried to completion.

#### iv. Liquid chromatography and mass spectrometry

Single-shot LC-MS/MS analyses were performed on 50% of each sample. The remaining 50% of each sample was fractionated using bRP StageTip fractionation. For single shot and all fractionated samples, desalted peptides were resuspended in 9 μL of 3% MeCN/0.1% FA and 4 μL was injected. For serum samples, an Orbitrap Fusion Lumos Tribrid Mass Spectrometer (ThermoFisher Scientific) was used. For all other plexes, an Orbitrap Exploris 480 (ThermoFisher Scientific) was used. Mass spectrometers were coupled online to a Proxeon Easy-nLC 1200 (ThermoFisher Scientific) as previously described56. Briefly, 4 μL of each sample was loaded at onto a microcapillary column (360 μm outer diameter × 75 μm inner diameter) containing an integrated electrospray emitter tip (10 μm), packed to approximately 24 cm with ReproSil-Pur C18-AQ 1.9 μm beads (Dr. Maisch GmbH) and heated to 50 °C. bRP fractionated samples were analyzed using a 110 min LC–MS. Mobile phase flow rate was 200 nL/min, comprises 3% acetonitrile/0.1% formic acid (Solvent A) and 90% acetonitrile /0.1% formic acid (Solvent B). The 110-min LC– MS/MS method used the following gradient profile: (min:%B) 0:2; 1:6; 85:30; 94:60; 95:90; 100:90; 101:50; 110:50 (the last two steps at 500 nL/min flow rate).. Data acquisition was done in the data-dependent mode acquiring HCD MS/MS scans (r = 15,000) after each MS1 scan (r = 60,000) on the top 12 most abundant ions using an MS1 AGC target of 4 x 105 and an MS2 AGC target of 5 × 10^4^. The maximum ion time utilized for MS/MS scans was 120 ms; the HCD-normalized collision energy was set to 36 (Fusion Lumos) or 28 (Exploris 480); the dynamic exclusion time was set to 20 s, and the peptide match and isotope exclusion functions were enabled. Charge exclusion was enabled for charge states that were unassigned, 1 and >7.

### MS data analysis

All protein trafficking MS data were analyzed using Spectrum Mill software package v 7.07 (proteomics.broadinstitute.org)). Similar MS/MS spectra acquired on the same precursor m/z within ±60 s were merged. MS/MS spectra were excluded from searching if they were not within the precursor MH+ range of 600–6000 Da or if they failed the quality filter by not having a sequence tag length >0. MS/MS spectra were searched against a UniProt mouse database with a release date of December 28, 2017 containing 46,519 proteins and 264 common contaminants modified to include GFP, mKate2 and BirA*G3-ER. All spectra were allowed ±20 ppm mass tolerance for precursor and product ions, 40% minimum matched peak intensity, and “trypsin allow P” enzyme specificity with up to 2 missed cleavages. The fixed modifications were carbamidomethylation at cysteine, and TMT6 at N-termini. The variable modifications used were oxidized methionine and N-terminal protein acetylation. Individual spectra were automatically designated as confidently assigned using the Spectrum Mill autovalidation module. Specifically, a target-decoy-based false-discovery rate (FDR) scoring threshold criteria via a two-step auto threshold strategy at the spectral and protein levels was used. First, peptide mode was set to allow automatic variable range precursor mass filtering with score thresholds optimized to yield a spectral level FDR of <1.2%. A protein polishing autovalidation was applied to further filter the peptide spectrum matches using a target protein level FDR threshold of 0. Following autovalidation, a protein–protein comparison table was generated, which contained experimental over control TMT ratios. For all experiments, non-mouse contaminants and reverse hits were removed. Furthermore, the data were median normalized. For serum data, we performed a moderated T-test (limma R package v4.1) to identify proteins significantly enriched in the experimental conditions compared to controls. We corrected for multiple hypotheses (Benjamini–Hochberg procedure). Any protein with an adjusted p-value of less than 0.05 and a log2 fold change greater than 1 was considered statistically enriched. For tissue data, we used ES method described below (MS hit analysis) to identify enriched proteins.

### MS Hit Analysis

To identify enriched proteins from MS data, we established threshold TMT ratios for hit-calling using positive control (PC) and negative control (NC) protein lists. For PC list, we used UniProt annotated secreted proteins, while the NC list was the UniProt overlapping list of transcription factors and nuclear proteins, and cytoskeletal genes. Note that the NC list was compared with secreted, receptors, ER proteins, and overlapping genes were removed.

Proteins identified by MS were compared to the PC and NC lists and assigned to as being a PC or NC protein. For each experiment (liver, brain, and kidney), there were 9 TMT ratio comparisons: we calculate the TMT ratios of every *Sox2-BirA*G3* sample over every control samples (*Sox2-BirA*G3*-1/ control-1, *Sox2-BirA*G3*-1/ control-2, *Sox2-BirA*G3*-1/ control-3, *Sox2-BirA*G3*-2/ control-1, *Sox2-BirA*G3*-2/ control-2, *Sox2-BirA*G3*-2/ control-3, *Sox2-BirA*G3*-3/ control-1, *Sox2-BirA*G3*-3/ control-2, and *Sox2-BirA*G3*-3/ control-3). The false positive rate (FPR) is calculated using the equation below^25^,

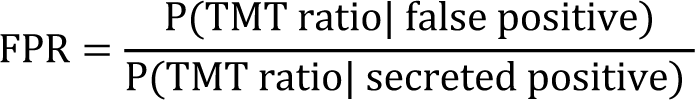

The denominator is the conditional probability of finding a known secreted protein in this range, which is calculated as the percentage of proteins on the PC list in this range over all proteins identified on the PC list. The numerator is the conditional probability of finding a false positive protein in a particular TMT ratio range. The result calculated using this equation represents the percentage of false positive proteins in this TMT ratio range over the total false positive proteins identified. We plotted FPR over TMT ratio range (Supplementary Fig. 4E; other plots available upon request) and selected the TMT ratio cutoff based on an FPR of 0.1 (Supplementary Fig. 4D; other plots available upon request), which means that a protein is 10 times more likely to be a true secreted protein than a false positive. Enrichment score (ES) of a specific protein was defined as the number of TMT ratios that exceeds the TMT ratio cutoff. Thus, proteins with ES of 0 are background, proteins with an ES of 1 are lower confidence hits, and proteins with ES of 9 are highest confidence hits.

### *Alb-BirA*G3* Serum Analysis by MS (corresponding to **Figure 5-6** and Extended Figures 10)

After streptavidin beads pulldowns, *Alb-BirA*G3* and control samples were sent to the UCLA proteomics core, Department of Biological Chemistry, Geffen School of Medicine at UCLA for MS.

#### i. Serum Sample Digestion

Streptavidin-bound proteins were reduced and alkylated on bead via sequential 20-minute incubations with 5mM TCEP and 10mM iodoacetamide at room temperature in the dark while being mixed at 1200 rpm in an Eppendorf thermomixer. Proteins were then digested by the addition of 0.1μg Lys-C (FUJIFILM Wako Pure Chemical Corporation, 125-05061) and 0.8μg Trypsin (Thermo Scientific, 90057) while shaking at 37°C overnight.

#### ii. TMT Labeling and CIF Fractionation

The supernatant was transferred to new tubes and 8 μl of carboxylate-modified magnetic beads (CMMB, and also widely known as SP3^60^) was added to each sample. 100% acetonitrile was added to each sample to increase the final acetonitrile concentration to >95% and induce peptide binding to CMMB. CMMB were then washed 3 times with 100% acetonitrile and then resuspended with TMT labeling buffer. 25 ug of each sample was labeled using TMT10 plex Isobaric Labels (Thermo Fisher Scientific) and the resulting 8 labeled samples were pooled. The pooled sample was fractionated by CMMB-based Isopropanol Gradient Peptide Fractionation (CIF) method^61^ into 3 fractions before MS analysis.

#### iii. LC-MS Acquisition and Analysis

Fractionated samples were separated on a 75uM ID x 25cm C18 column packed with 1.9μm C18 particles (Dr. Maisch GmbH) using a 140-minute gradient of increasing acetonitrile and eluted directly into a Thermo Orbitrap Fusion Lumos mass spectrometer where MS spectra were acquired using SPS-MS3.

Protein identification was performed using MaxQuant^62^ v 1.6.17.0. The complete Uniprot mouse proteome reference database (UP000000589) was searched for matching MS/MS spectra. Searches were performed using a 20 ppm precursor ion tolerance. TMT10plex was set as a static modification on lysine and peptide N terminal. Carbamidomethylation of cysteine was set as static modification, while oxidation of methionine residues and N-terminal protein acetylation were set as variable modifications. LysC and Trypsin were selected as enzyme specificity with maximum of two missed cleavages allowed. 1% false discovery rate was used as a filter at both protein and PSM levels.

Statistical analysis was conducted with the MSstatsTMT Bioconductor package^63^. The abundance of proteins missing from one condition but found in more than 2 biological replicates of the other condition for any given comparison were estimated by imputing intensity values from the lowest observed MS1-intensity across samples and p-values were randomly assigned to those between 0.05 and 0.01 for illustration purposes.

### Data analysis and statistics

Data was analyzed using Microsoft Excel, R (version 4.0.0 (2020-04-24), Platform: x86_64-appledarwin17.0 (64-bit); RStudio Version 1.3.959) and Python. For secretion annotations, proteins were annotated based on the subcellular localization data from UniProt and the cellular component data from National Center for Biotechnology Information (NCBI) (https://www.ncbi.nlm.nih.gov). Proteins in fasta formats were uploaded to SignalP5.0 (https://services.healthtech.dtu.dk/service.php?SignalP-5.0) and TMHMM (v 2.0) (https://services.healthtech.dtu.dk/service.php?TMHMM-2.0) for the prediction of SignalP and transmembrane helix, separately. EnhancedVolcano was used to generate volcano plots based on log_2_FC and p value. We ran TissueEnrich (https://bioconductor.org/packages/TissueEnrich)^32^ on a list of proteins to look for enrichment for tissue-specific genes using mouse ENCODE datasets. Gene ontology function annotation was performed on two platforms-DAVID (https://david.ncifcrf.gov) and EnrichGO in clusterProfiler (3.16.1). The top GO terms were visualized with dotplot in ggplot2. PCA was used to study the similarities between samples. The analysis was conducted without filtering any proteins. For ectodomain shedding analysis, we wrote a python program to map all the peptides identified in mass spec for each protein to their corresponding full-length protein. The reference sequences were annotated based on UniProt topology information. Domain information was based on SMART (http://smart.embl-heidelberg.de).

### Data Availability

The original mass spectra and the protein sequence databases used for searches have been deposited in the public proteomics repository MassIVE (htt://massive.ucsd.edu) and are accessible at ftp://MSV000088848@massive.ucsd.edu. RNA sequencing data is available under BioProject PRJNA808087 from NCBI SRA at https://www.ncbi.nlm.nih.gov/sra. The following public databases were used: Uniprot (https://www.uniprot.org), mouse (https://www.uniprot.org/proteomes/UP000000589), SignalP 5.0 (https://services.healthtech.dtu.dk/service.php?SignalP-5.0), TMHMM 2.0 (https://services.healthtech.dtu.dk/service.php?TMHMM-2.0), Corresponding authors will provide original data upon request. Source data are provided with this paper.

---

**Extended Fig. 1.**
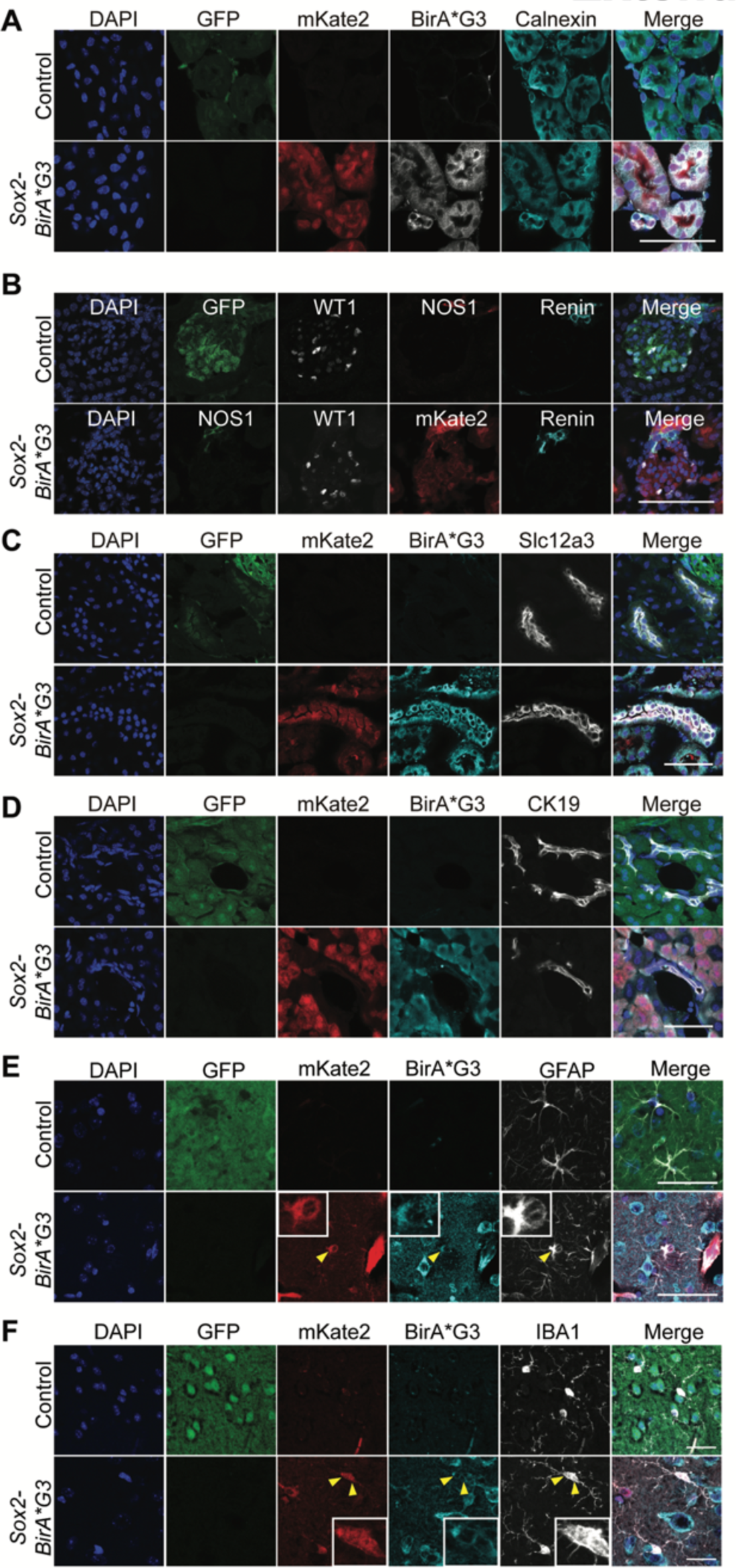
BirA*G3 expression and colocalization with ER marker Calnexin in *Sox2-BirA*G3* mice. (A) Immunofluorescence staining shows expression of native GFP, native mKate2, BirA*G3, and ER marker Calnexin in *Sox2-BirA*G3* and control mouse kidney. Scale Bar: 50μm. (B) Immunofluorescence staining shows expression of native GFP, native mKate2, BirA*G3, Nitric Oxide Synthase 1 (NOS1), Wilms’ tumor 1 (WT1) and Renin in the kidney sections of *Sox2-BirA*G3* and control mice. Scale Bar: 50μm. (C) Immunofluorescence staining shows expression of native GFP, native mKate2, BirA*G3, and Solute Carrier Family 12 Member 3 (Slc12a3), which marks distal convoluted tubule, in the kidney sections of *Sox2-BirA*G3* and control mice. Scale Bar: 50μm. (D) Immunofluorescence staining shows expression of native GFP, native mKate2, BirA*G3, and cytokeratin (CK19), a marker for bile duct, in the liver sections of *Sox2-BirA*G3* and control mice. Scale Bar: 50μm. (E) Immunofluorescence staining shows expression of native GFP, native mKate2, BirA*G3, and GFAP, a marker for glia, in the brain sections of *Sox2-BirA*G3* and control mice. Cells indicated by yellow arrow are magnified as in the rectangles. Scale Bar: 50μm. (F) Immunofluorescence staining shows expression of native GFP, native mKate2, BirA*G3, and IBA-1, a marker for microglia, in the brain sections of *Sox2-BirA*G3* and control mice. Cells indicated by yellow arrow are magnified as in the rectangles. Scale Bar: 25μm.

**Extended Fig. 2.**
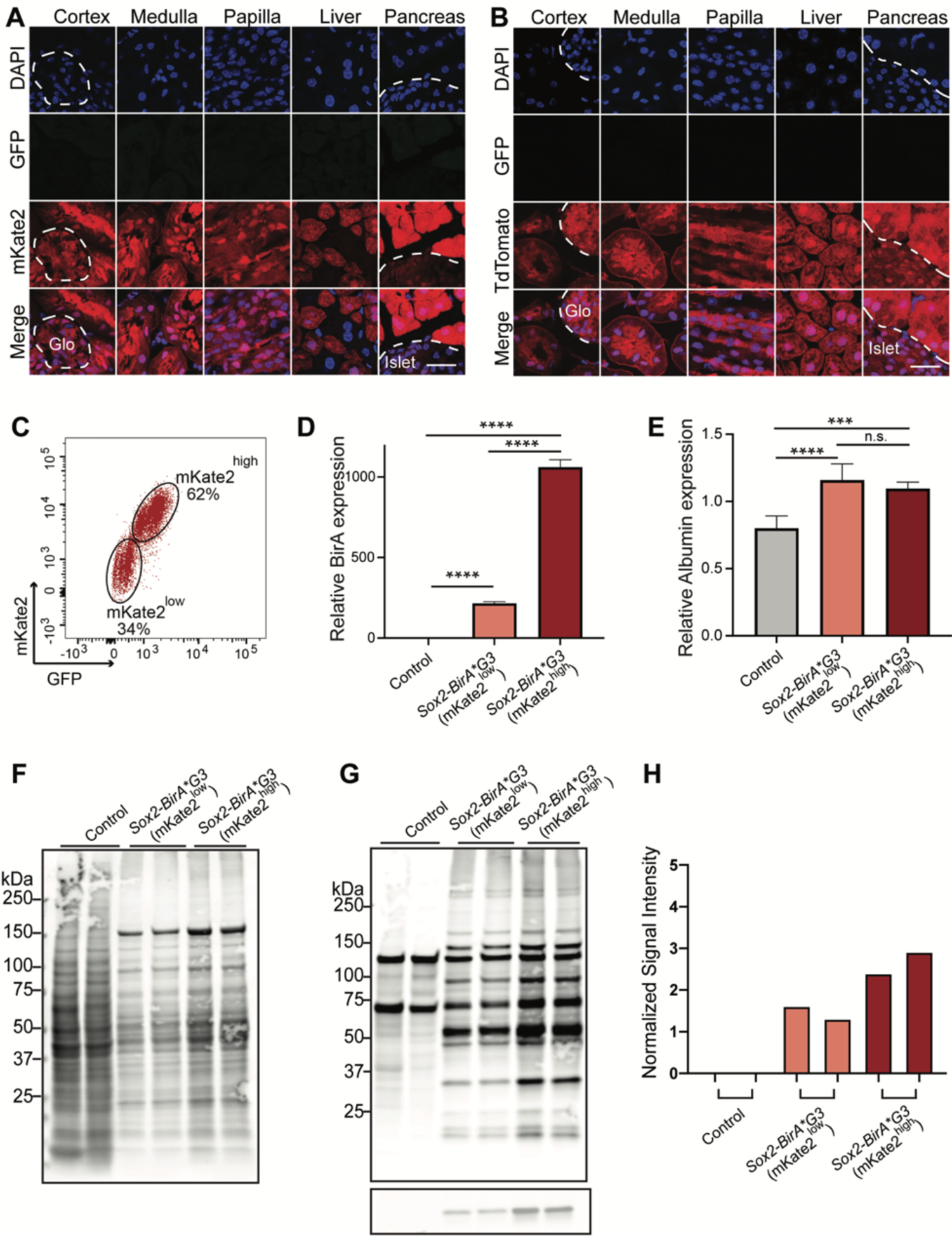
Partial BirA silencing observed in *Sox2-BirA*G3* mice was affirmed by comparing with *Sox2-Cre; TdTomato* mice. (A) Representative high magnification images of native fluorescence expression in *Sox2-BirA*G3* kidney, livers, and pancreas. Scale Bar: 25μm. (B) Representative high magnification images of native fluorescence expression in *Sox2-TdTomato* kidney, liver, and pancreas. Scale Bar: 25μm. (C) mKate2^low^ and mKate2^high^ hepatocytes were sorted from *Sox2-BirA*G3* liver by flow cytometry based on native GFP and native mKate2 expression. (D) BirA*G3 RNA expression was analyzed by Q-PCR in mKate2^low^ and mKate2^high^ hepatocytes of *Sox2-BirA*G3* liver compared to total control liver. One-way ANOVA with Tukey’s multiple comparisons test were used to test for significant differences between individual groups. (E) Albumin RNA expression was analyzed by Q-PCR in mKate2^low^ and mKate2^high^ hepatocytes of *Sox2-BirA*G3* liver compared to total control liver. One-way ANOVA with Tukey’s multiple comparisons test were used to test for significant differences between individual groups. (F) Total protein stain of total protein lysates from mKate2^low^ and mKate2^high^ hepatocytes of *Sox2-BirA*G3* liver compared to total control liver. Each lane is a biological replicate from individual mice (n=2/genotype). (G) Western blotting of total protein lysates from mKate2^low^ and mKate2^high^ hepatocytes of *Sox2-BirA*G3* liver compared to total control liver. Upper: Streptavidin labeling. Lower: BirA*G3 (∼35kDa). Each lane is a biological replicate from individual mice (n=2/genotype). (H) Western blot quantification of BirA*G3 levels in mKate2^low^ and mKate2^high^ hepatocytes of *Sox2-BirA*G3* liver compared to total control liver. Each lane is a biological replicate from individual mice (n=2/genotype).

**Extended Fig. 3.**
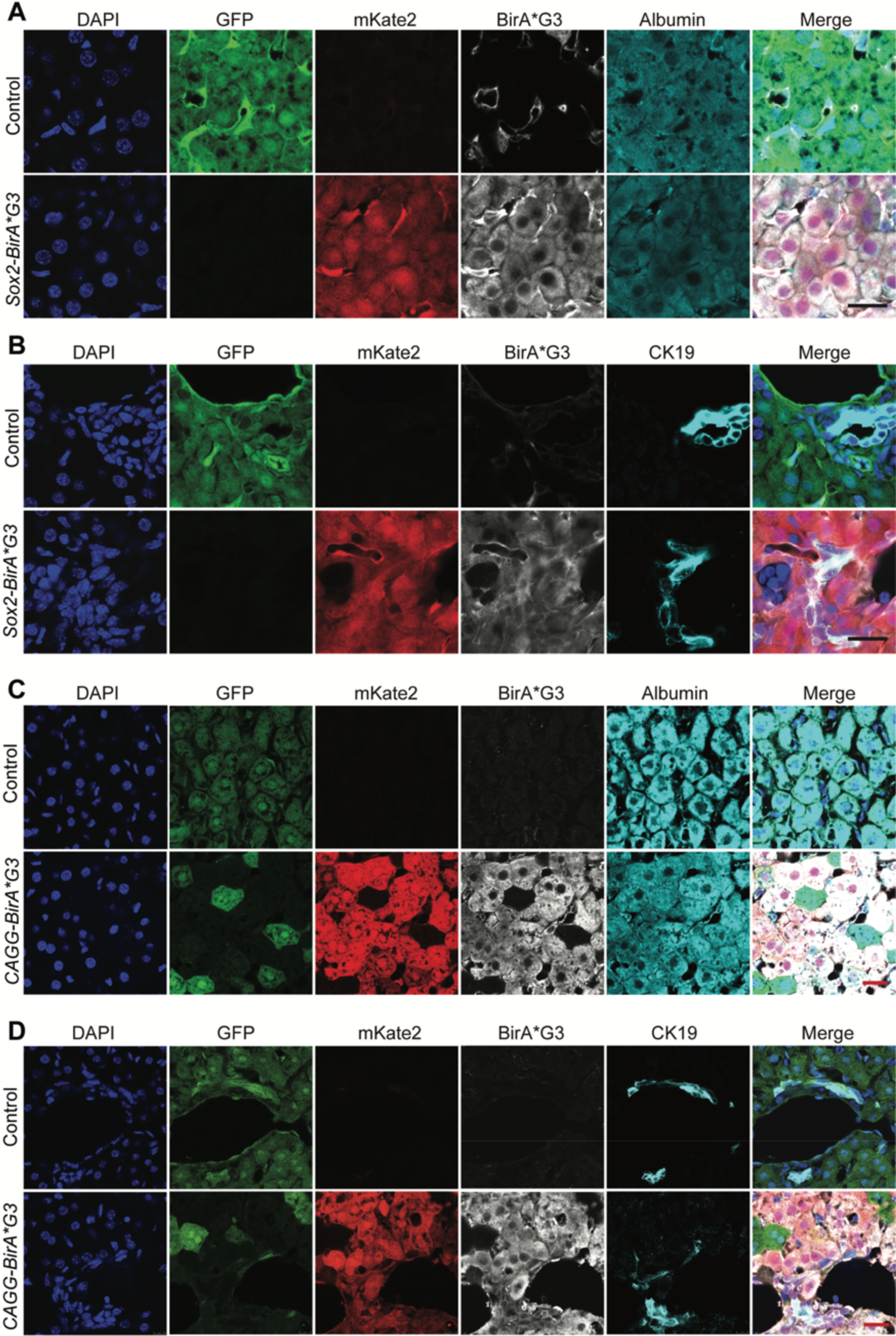
Homogenous mKate2 and BirA*G3 expression was detected in *Sox2-BirA*G3* and control mouse pups at 10 days post birth and in CRE-induced liver cells from *CAGG-BirA*G3* mice. (A) Immunofluorescence images showed native GFP, native mKate2, BirA*G3, and Albumin in the liver sections of *Sox2-BirA*G3* and control mice at 10 days post birth. Scale bar: 20μm. (B) Immunofluorescence images showed native GFP, native mKate2, BirA*G3, and CK19 in the liver sections of *Sox2-BirA*G3* and control mice at 10 days post birth. Scale bar: 20μm. (C) Immunofluorescence images showed native GFP, native mKate2, BirA*G3, and Albumin in the liver sections of *CAGG-BirA*G3* and control mice. Scale bar: 20μm. (D) Immunofluorescence images showed native GFP, native mKate2, BirA*G3, and CK19 in the liver sections of *CAGG-BirA*G3* and control mice. Scale bar: 20μm.

**Extended Fig. 4.**
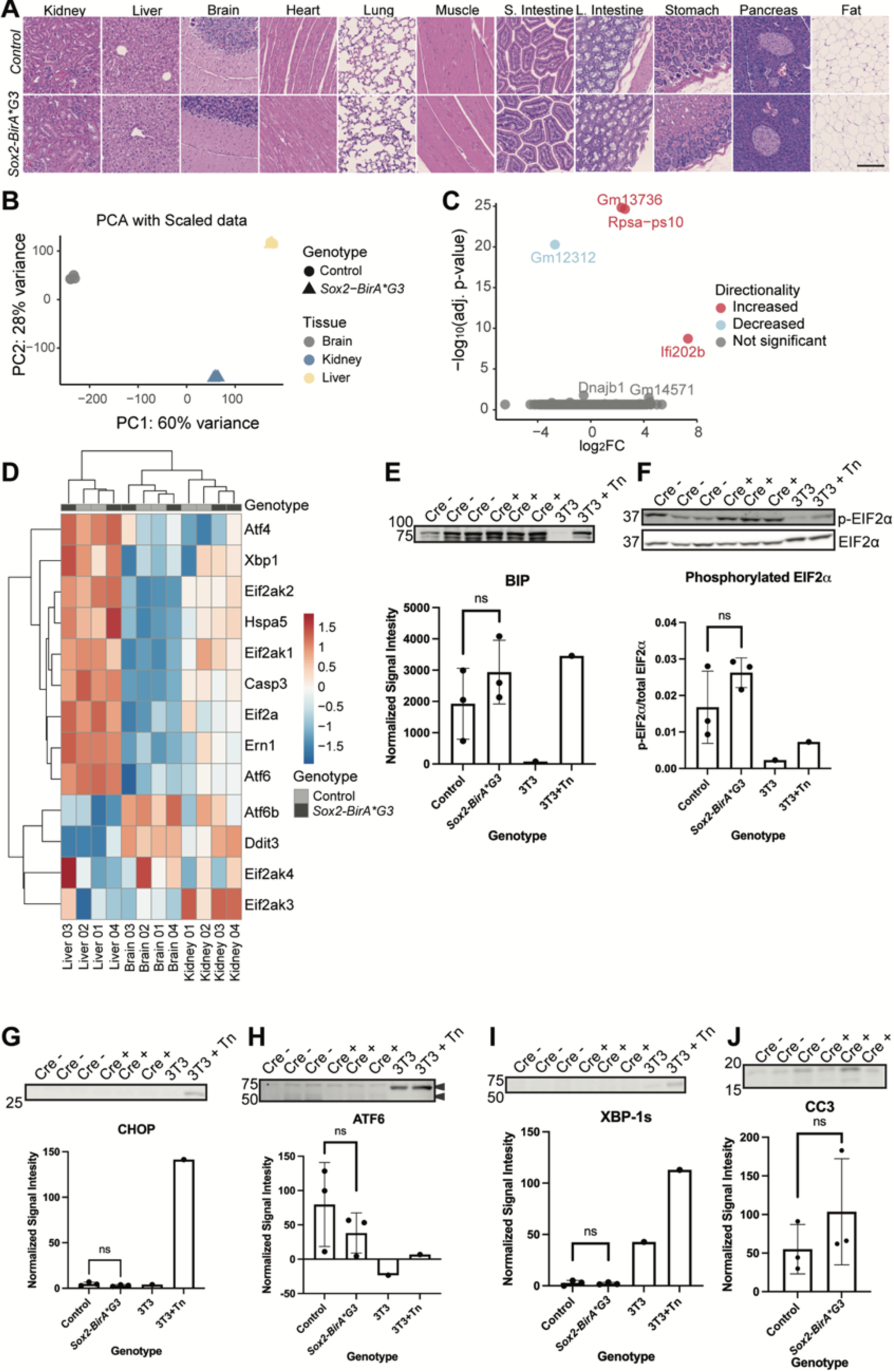
No detrimental effects were observed in *Sox2-BirA*G3* and control mice. (A) Representative images of hematoxylin and eosin staining of paraffin-sectioned tissues from *Sox2-BirA*G3* and control mice with 7-day biotin chow. Scale Bar: 100μm. (B) PCA of *Sox2-BirA*G3* and control RNA sequencing samples from all tissues (liver, brain, and kidney). Each point represents one sample (biological replicate) where color is by tissue type and shape is by genotype. (C) Volcano plot showing all differentially expressed genes (DEGs) in *Sox2-BirA*G3* samples compared to control samples for all three tissues. Increased (red) are DEGs in the *Sox2-BirA*G3* condition and decreased (blue) are DEGs in the control condition. (D) Heatmap of ER stress and unfolded protein response genes log_2_ fold change (scale) from differential expression analysis between Sox*2-BirA*G3* and control samples showing hierarchical clustering by tissue. (E-J) Western blot and quantification normalized to total protein stain (not shown) except EIF2α (normalized to total EIF2α) of ER stress markers BIP (Grp78) (E, p-value 0.32), phosphorylated (p) EIF2α and total EIF2α (F, p-value 0.23), CHOP (G, p-value 0.26), ATF6 (H, p-value 0.37), XBP-1s (I, p-value 0.75), and apoptotic marker CC3 (J, p-value 0.35) of 25μg liver lysate from control and *Sox2-BirA*G3* samples compared to ER stress controls (mouse 3T3 cells treated with or without Tunicamycin (Tn)). Arrows in D indicate molecular weights for total ATF6 (predicted 75 kDa) and cleaved/active ATF6 (50 kDa). Each lane is a biological replicate from individual mice (n=3/genotype). Statistical significance (significance = p-value < 0.05; ns: non-significant) was calculated using a two tailed, Welch’s t-test between *Sox2-BirA*G3* and control samples.

**Extended Fig. 5.**
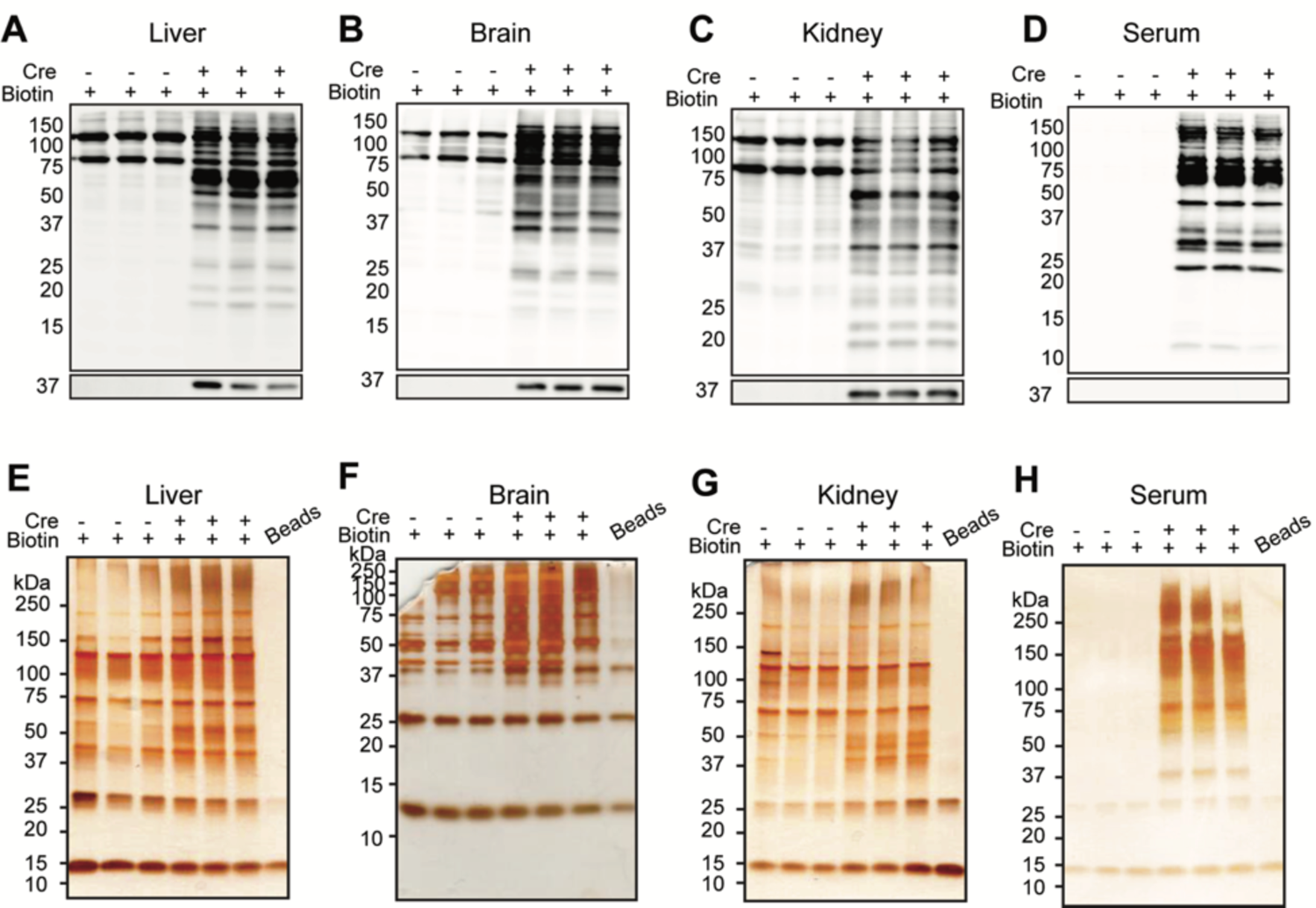
Streptavidin purification of biotinylated proteins for LC-MS/MS from *Sox2-BirA*G3 mice*. (A-D) Western blotting of streptavidin affinity purified biotinylated proteins from liver (A), brain (B), kidney (C), serum (D) in *Sox2-BirA*G3* mice compared to control mice. Upper: Streptavidin labeling. Lower: BirA*G3 (∼35kDa). Each lane is a biological replicate from individual mice (n=3/genotype). (E-H) Silver stain of streptavidin affinity purified biotinylated proteins from liver (E), brain (F), kidney (G), and serum (H) in *Sox2-BirA*G3* mice compared to control mice. Bead lanes are affinity purification negative control without protein input to show streptavidin contribution to bound fractions from beads. Each lane is a biological replicate from individual mice (n=3/genotype).

**Extended Fig. 6.**
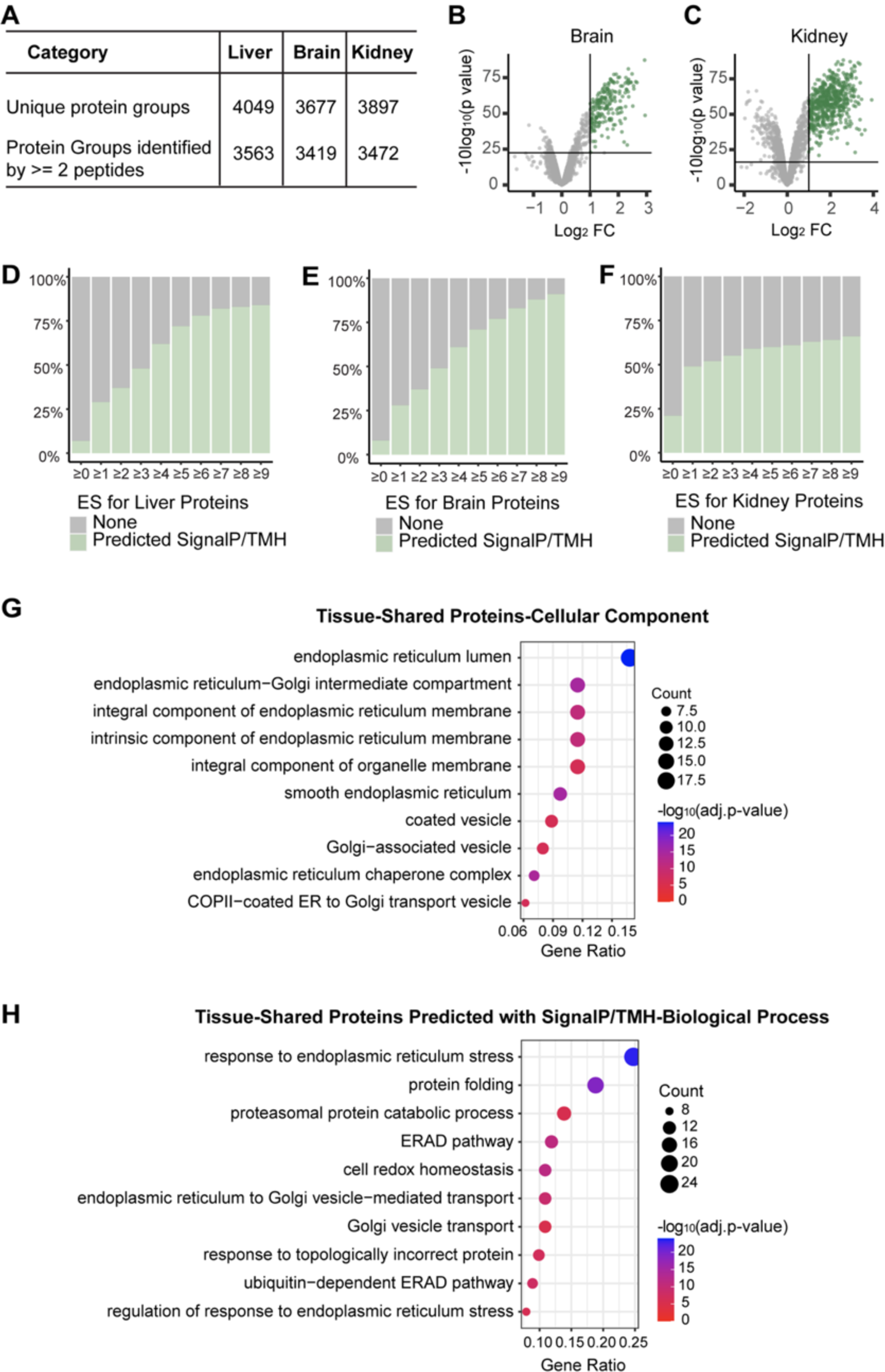
Quantitative LC-MS/MS data analysis for three tissues. (A) Summary of information obtained from quantitative LC-MS/MS. (B-C) Volcano plots of proteins detected in brain (B) and kidney (C) of *Sox2-BirA*G3* mice compared to control mice after streptavidin pulldown. Log_2_ FC were plotted on the x-axis and −10log_10_ (p value) were plotted on the y-axis. Significantly enriched proteins (adj. p-value< 0.05 and log_2_FC>1) in *Sox2-BirA*G3* mice compared to control mice are shown in green or red. (D-F) Percentage of proteins with predicted SignalP/TMH in each ES category. As the ES increases, the fraction of proteins with predicted SignalP or TMH increases. (G) Shared enriched proteins among three tissues (113 proteins) were analyzed with clusterProfiler (3.16.1) EnrichGO analysis for cellular components annotation. Gene ratio indicates the percentage of genes annotated with the term over the total number of genes in the list. (H) Shared enriched proteins among three tissues predicted with SignalP/TMH were analyzed with clusterProfiler (3.16.1) EnrichGO analysis for biological process annotation. Gene ratio indicates the percentage of genes annotated with the term over the total number of genes in the list.

**Extended Fig. 7.**
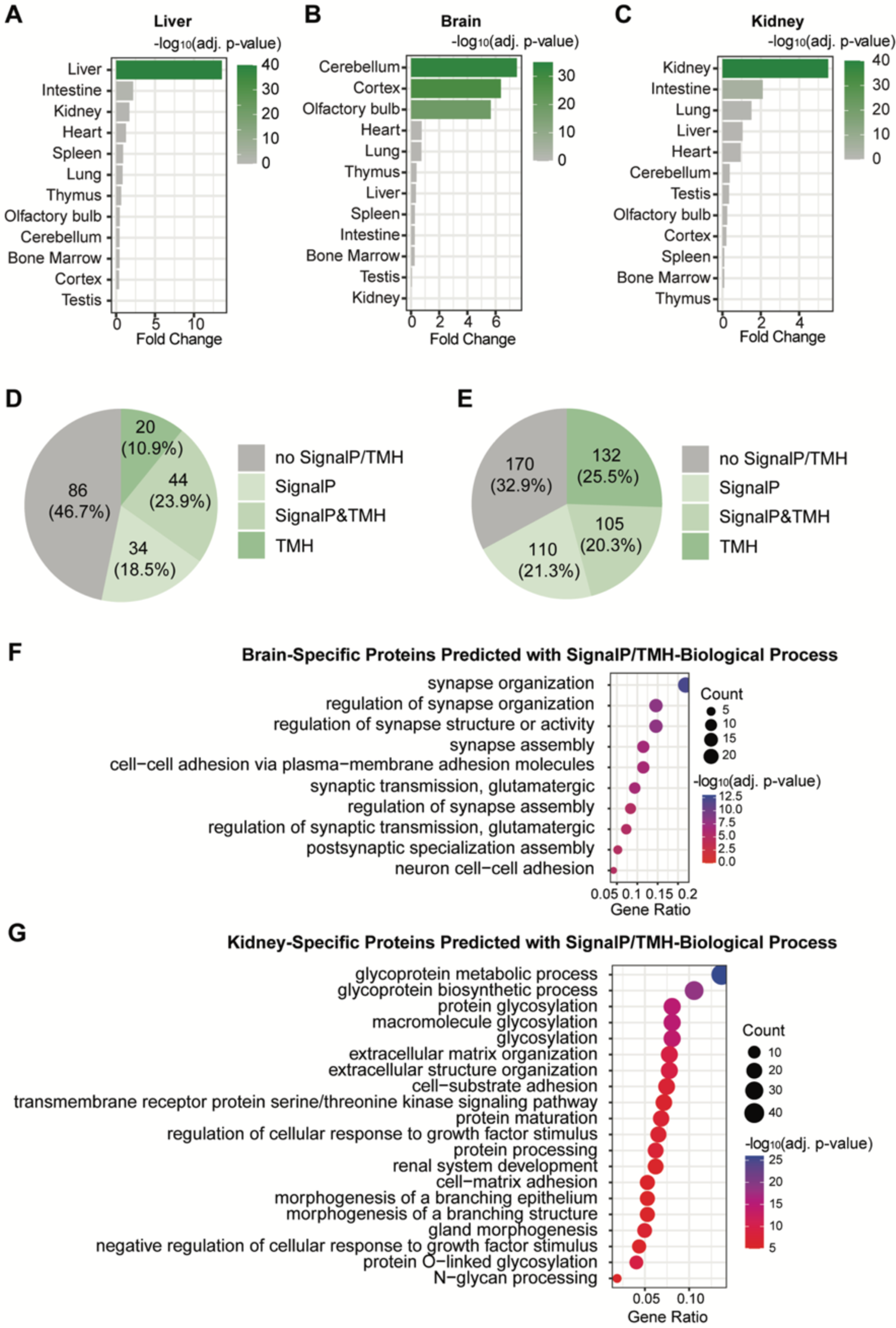
Tissue-specific analysis for tissue-specific enriched proteins. (A-C) Tissue-specific enriched proteins from liver (A), brain (B), and kidney (C) were analyzed with TissueEnrich, a tool that calculates tissue-specific gene enrichment in an input gene set. (D-E) Pie plots showed the number of brain-specific enriched proteins (184 proteins) (D) and kidney-specific enriched proteins (517 proteins) (E) predicted with SignalP/TMH. (F) Brain-specific enriched proteins predicted with SignalP/TMH were analyzed with clusterProfiler (3.16.1) EnrichGO analysis for biological process. Gene ratio indicates the percentage of genes annotated with the term over the total number of genes in the list. (G) Kidney-specific enriched proteins predicted with SignalP/TMH were analyzed with clusterProfiler (3.16.1) EnrichGO analysis for biological process. Gene ratio indicates the percentage of genes annotated with the term over the total number of genes in the list.

**Extended Fig. 8.**
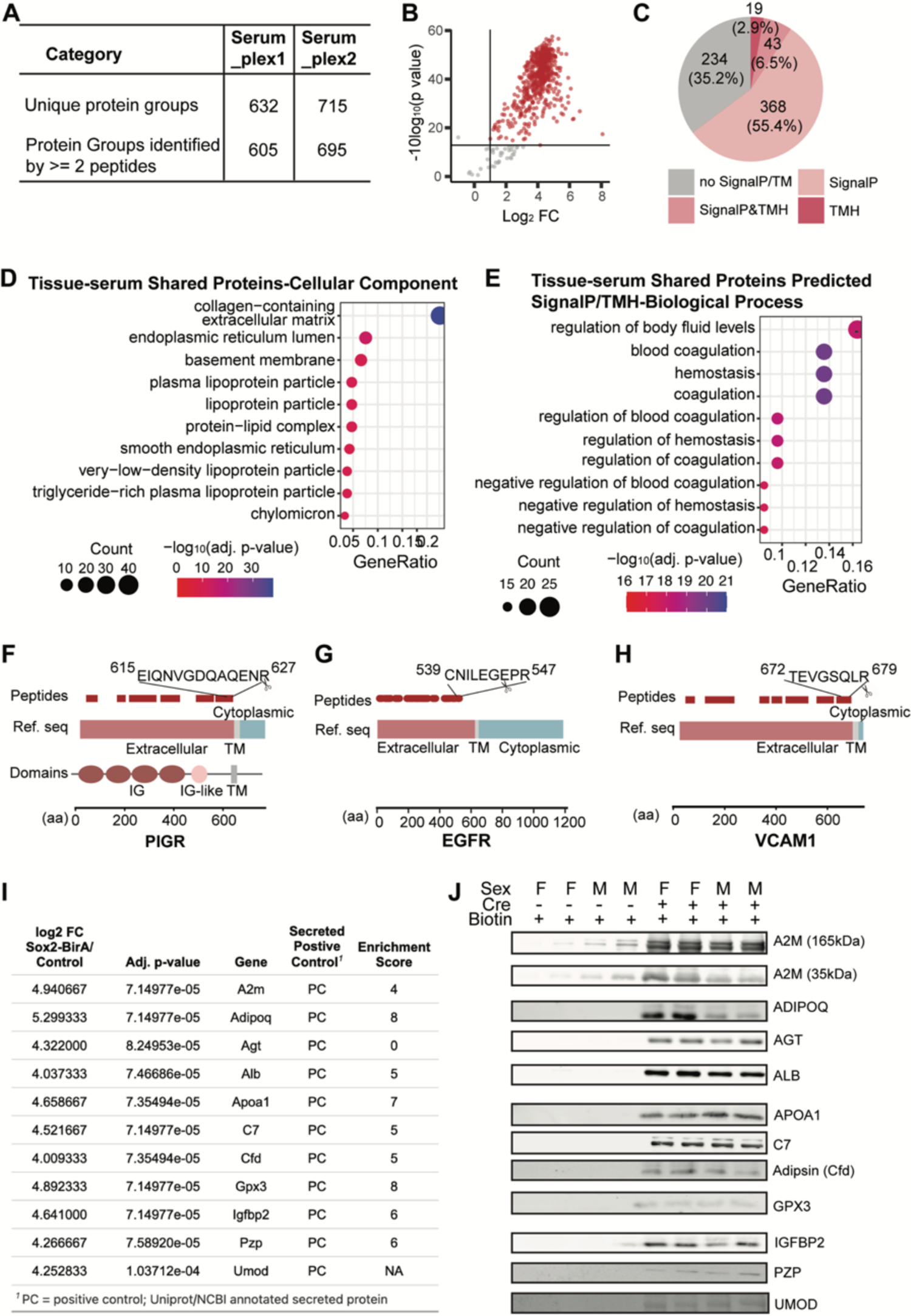
Quantitative MS data analysis and western blot validation for serum. (A) Summary of information obtained from quantitative MS. (B) Volcano plo of proteins detected in serum (plex1) of *Sox2-BirA*G3* mice compared to control mice after streptavidin pulldown. Log_2_ FC were plotted on the x-axis and −10Log_10_ (p value) were plotted on the y-axis. Significantly enriched proteins (adj. p-value< 0.05 and log_2_FC>1.0) in *Sox2-BirA*G3* mice compared to control mice are shown in red. Volcano plot of proteins detected in serum (plex2) of *Sox2-BirA*G3* mice compared to control mice after streptavidin pulldown were not shown. (C) Pie chart displayed the distribution of serum enriched proteins with predicted SignalP/TMH. (D) Shared enriched proteins between serum and three tissues were analyzed with clusterProfiler (3.16.1) EnrichGO enrichment analysis for cellular component annotation. (E) Shared enriched proteins between serum and three tissues were analyzed with clusterProfiler (3.16.1) EnrichGO enrichment analysis for biological process annotation. (F) Schematic of detected peptides for PIGR mapped onto its respective reference sequences with SMART protein domain annotation. Reference sequence is annotated with extracellular, TM and cytoplasmic based on UniProt topology information. Amino acid sequences of the most C-terminal peptide are labeled. IG: immunoglobulin. (G) Schematic of detected peptides for EGFR mapped onto its respective reference sequences. Reference sequence is annotated with extracellular, TM and cytoplasmic based on UniProt topology information. Amino acid sequences of the most C-terminal peptide are labeled. (H) Schematic of detected peptides for VCAM1 mapped onto its respective reference sequences. Reference sequence is annotated with extracellular, TM and cytoplasmic based on UniProt topology information. Amino acid sequences of the most C-terminal peptide are labeled. (I) Mass spectrometry hits of well characterized secreted proteins showing Log_2_FC and significant adjusted p-value enrichment method compared to enrichment score (ES) method. (J) Western blot validation of streptavidin affinity purified serum hits from (I) in additional control and *Sox2-BirA*G3* mice (n=2/sex). Input: 100, 200μg, or 600μg protein. Each lane is a biological replicate from individual mice (n=4/genotype).

**Extended Fig. 9.**
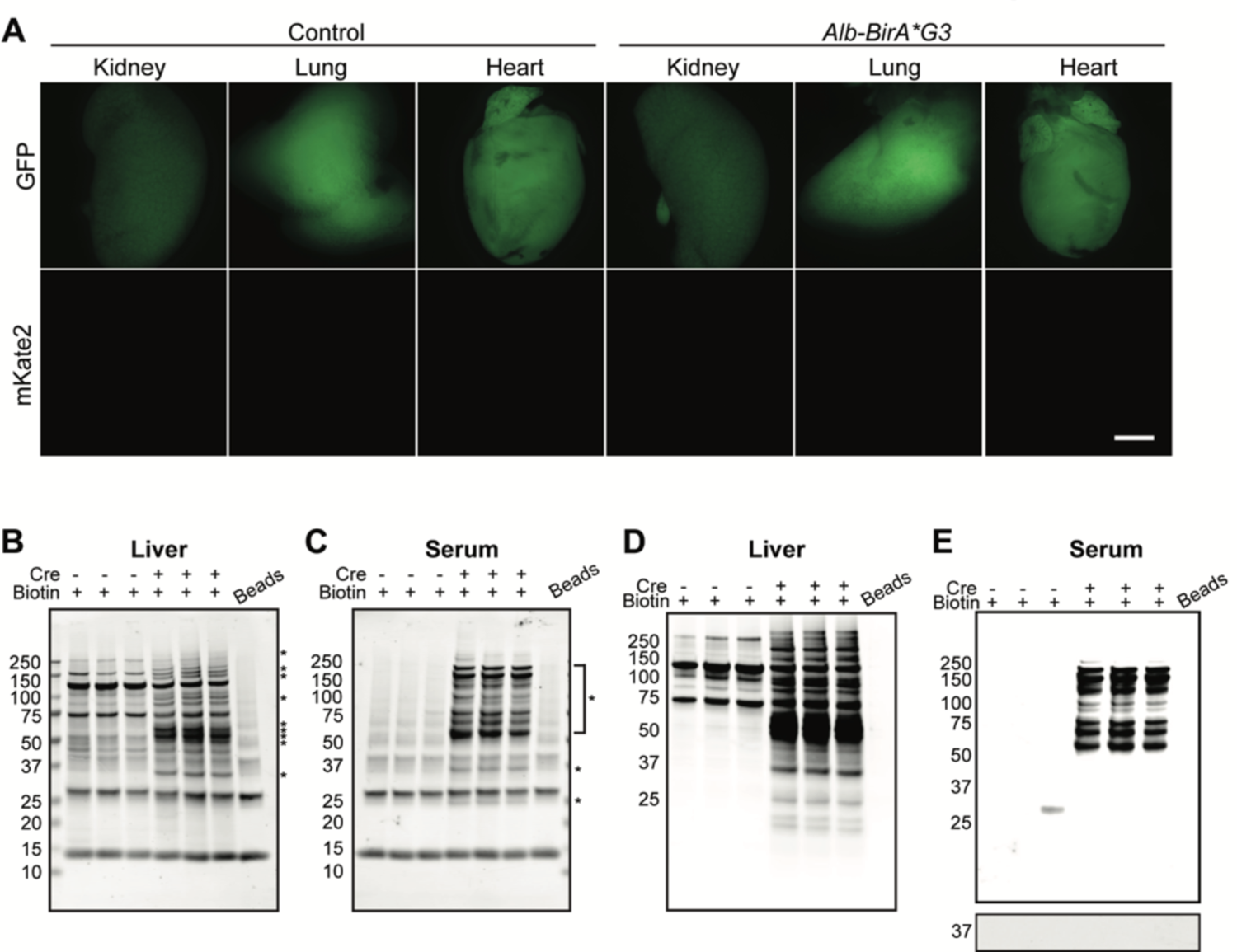
Characterization of *Alb-BirA*G3* and control mice. (A) Bright field images of well-perfused organs from *Alb-BirA*G3* and control mice in comparison with organs without perfusion. (B-C) Total protein stain of affinity purified biotinylated proteins from liver (B) and serum (C) in *Alb-BirA*G3* mice compared to control mice. Specific bands are indicated by asterisks. Bead lanes are affinity purification negative control without protein input to show streptavidin contribution to bound fractions from beads. Each lane is a biological replicate from individual mice (n=3/genotype). (D-E) Streptavidin labeling of affinity purified biotinylated proteins from liver (D) and serum (E) in *Alb-BirA*G3* mice compared to control mice. Lower: BirA*G3 (∼35kDa). Bead lanes are affinity purification negative control without protein input. Each lane is a biological replicate from individual mice (n=3/genotype).

**Extended Figure 10.**
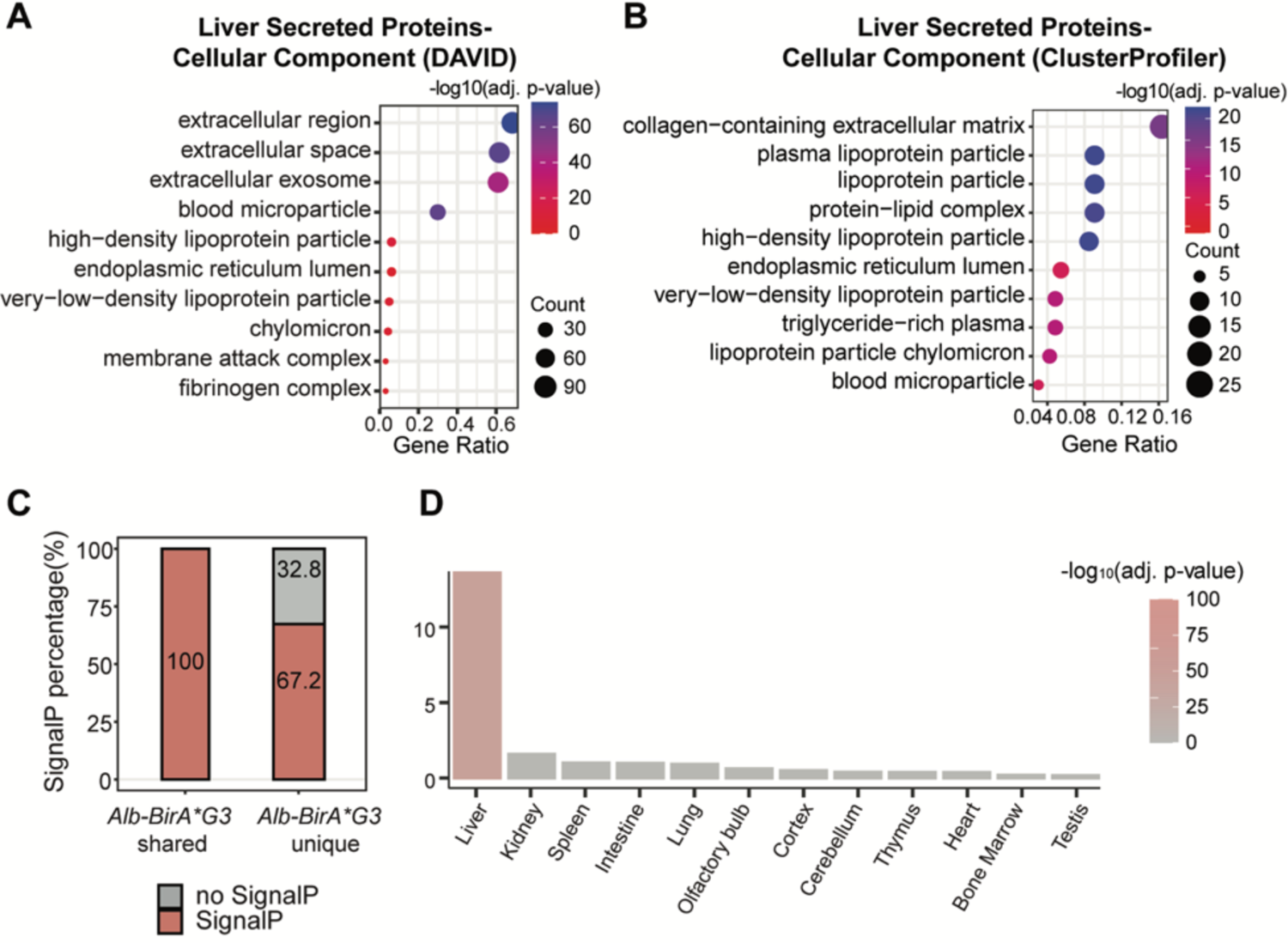
Analysis of *Alb-BirA*G3* and control serum enriched proteins. (A) Enriched serum proteins in *Alb-BirA*G3* mice were analyzed with DAVID analysis for cellular component annotation. Gene ratio indicates the percentage of genes annotated with the term over the total number of genes in the list. (B) Enriched serum proteins in *Alb-BirA*G3* mice were analyzed with clusterProfiler (3.16.1) EnrichGO enrichment analysis for cellular component annotation. Gene ratio indicates the percentage of genes annotated with the term over the total number of genes in the list. (C) Bar plot showed the percentage of unique (n=137) or shared (n=45) *Alb-BirA*G3* secreted proteins (compared with previous datasets) with predicted SignalP. (D) Unique *Alb-BirA*G3* secreted proteins (n=137) were analyzed with TissueEnrich to calculate tissue-specific gene enrichment.

